# ALTering cancer by triggering telomere replication stress through the stabilization of promoter G-quadruplex in *SMARCAL1*

**DOI:** 10.1101/2024.02.01.578345

**Authors:** Suman Panda, Tanaya Roychowdhury, Anindya Dutta, Sourio Chakraborty, Tanya Das, Subhrangsu Chatterjee

## Abstract

Most of the human cancers are dependent on telomerase to extend the telomeres. But ∼10% of all cancers use a telomerase-independent, homologous recombination mediated pathway called alternative lengthening of telomeres (ALT). Due to poor prognosis, the ALT status is not being considered yet in the diagnosis of cancer. No such specific treatment is available till date for ALT cancers. ALT positive cancers are dependent on replication stress to deploy DNA repair pathways to the telomeres to execute homology recombination mediated telomere extension. SMARCAL1 (SWI/SNF related, matrix-associated, actin-dependent regulator of chromatin, subfamily A-like 1) is associated with the ALT telomeres to resolve replication stress thus providing telomere stability. Thus, the dependency on replication stress regulatory factors like SMARCAL1 made it a suitable therapeutic target for the treatment of ALT positive cancers. In this study, we found a significant downregulation of SMARCAL1 expression by stabilizing the G-quadruplex (G4) motif found in the promoter of *SMARCAL1* by potent G4 stabilizers, like TMPyP4 and BRACO-19. SMARCAL1 downregulation led towards the increased localization of PML (promyelocytic leukaemia) bodies in ALT telomeres and triggered the formation of APBs (ALT-associated promyelocytic leukaemia bodies) in ALT positive cell lines, thus increasing telomere replication stress and DNA damage at genomic level. Induction of replication stress and hyper recombinogenic phenotype in ALT cells mediated by G4 stabilizing molecules already highlighted their possible application as a new therapeutic window to target ALT-positive tumors. In accordance with this, our study will also provide a valuable insight towards the development of G4 based ALT-therapeutics targeting *SMARCAL1*.

---

For telomere length maintenance, 90% of all the cancers are mainly dependent on the telomerase activity. But telomerase activity is completely absent in approximately 10% of all human cancers, where the telomere length maintenance mechanism is dependent on a homologous recombination mediated process called alternative lengthening of telomeres (ALT).^1 2^ ALT mechanism is commonly reported in several subtypes of sarcoma, glioblastoma and astrocytoma. Research already showed the nexus between the ALT status and prognosis in some tumor.^1^ The involvement of replication stress is severe in ALT positive cancers, which includes DNA damage at genomic level, unwanted chromosomal fusions, and genomic instability.^3 4^ ALT positive cells are highly sensitive towards further replicative impairment of the downstream replication stress response pathways.^5^ The replication stress response protein SMARCAL1 is a critical regulator of ALT activity in ALT positive cancer. SMARCAL1 is an ATP-dependent DNA annealing helicase, ubiquitously expressed in several human tissues, which is recruited to the stalled replication forks during replication stress, to restart replication through the remodeling of the surrounding environment.^6 7 8 9^

Recent study showed the accumulation of SMARCAL1 at ALT telomeres to resolve replication stress and found to restart stalled replication forks by triggering branch migration and fork regression, thus providing telomere stability.^10^ Thus, loss of SMARCAL1 leads towards DNA damage, changes in telomere heterogeneity, driving genomic instability and thus supporting a significant role of SMARCAL1 in the maintenance of ALT telomere stability. Recently, SMARCAL1 was found to be mutated in half of the ALT-positive glioblastoma samples, making it a potential therapeutic target.^11^ The strong scientific foundation already supported the induction of telomere-specific replication stress as a rapid and potent gateway to crush ALT positive cancer cells, thus making SMARCAL1 as a potential therapeutic target for the development of ALT specific treatment.^12 13 4^

In the past decades, non-canonical structure like G-quadruplex (G4) has emerged as a new intracellular therapeutic target due to its gene regulatory functions.^14 15 16 17 18 19^These G4s regulate the accuracy of gene expression and dictate the fate of several important biological events like telomere dysfunction, replication, oncogene transcriptions, genomic instability, ribosomal RNA biogenesis etc.^20 21 22^ Usually, these putative G4s act as a transcription repressor element to tune up the transcript threshold of the gene, thus regulating the cellular homeostasis. To understand the mechanism of transcription regulation of SMARCAL1 in ALT positive cancer cells, the promoter region of SMARCAL1 is extensively studied and we identified the presence of one putative G4 motif, which can dictate the G4 mediated fine tuning of SMARCAL1 expression. We also tried to understand the repercussion of stabilizing this G4 by widely reported G4 stabilizing molecules, i.e., TMPyP4 and BRACO-19. Overall, our study unravels the consequences of G4 stabilization in *SMARCAL1* promoter, resulting into the downregulation of SMARCAL1 and inducing global DNA damage, replication stress in the telomere (3’-TTAGGG) of ALT cancer cells, which can be used as a potent and expeditious route to destroy ALT cancer cells. Our study will surely open a new avenue towards the development of future ALT therapeutics by developing specific small molecules to stabilize G4 structure of *SMARCAL1*.

From bioinformatics studies using QGRS (G-quadruplex prediction software) server, one putative G4 motif in the sense strand of *SMARCAL1* P3 promoter (Chromosome 2, Gene ID: 50485; Ensembl: ENSG00000138375; NC_000002.12 GRCh38.p14 Primary Assembly), found from MatInspector was mapped. We named the putative G4 motif as P3 G4 wt (25 bases, -17 to +8) (Figure 1A). To evaluate our conviction of putative G4 motif formation, two oligonucleotides i.e., one wild type sequence (P3 G4 wt) and one mutated sequence (P3 G4 mut) were synthesized. Due to the dynamic nature of G4 formation, we mutated more number of Guanines to Thymine to reduce the probability of the mutated sequence to form other non-canonical structures. Then, CD spectra of these sequences were obtained in the wavelength range of 210–300 nm to investigate the formation of G4 structure and the respective topology. In CD spectra, P3 G4 wt sequence showed a characteristic positive peak at 260–265 nm and a negative dip around 240 nm, indicating an exclusive signature of parallel G4 topology (Figure S1).^23^ In comparison, the mutated sequence, P3 G4 mut did not evolve such characteristic peaks due to the absence of important guanine residues (Figure S2). Then, we calculated the melting temperature (Tm) of the putative G4 motif (Figure 1B). The putative G4 motif was found to be stabilized by K^+^ ions, which intercalate between consecutive G-tetrads.

**Figure 1.**
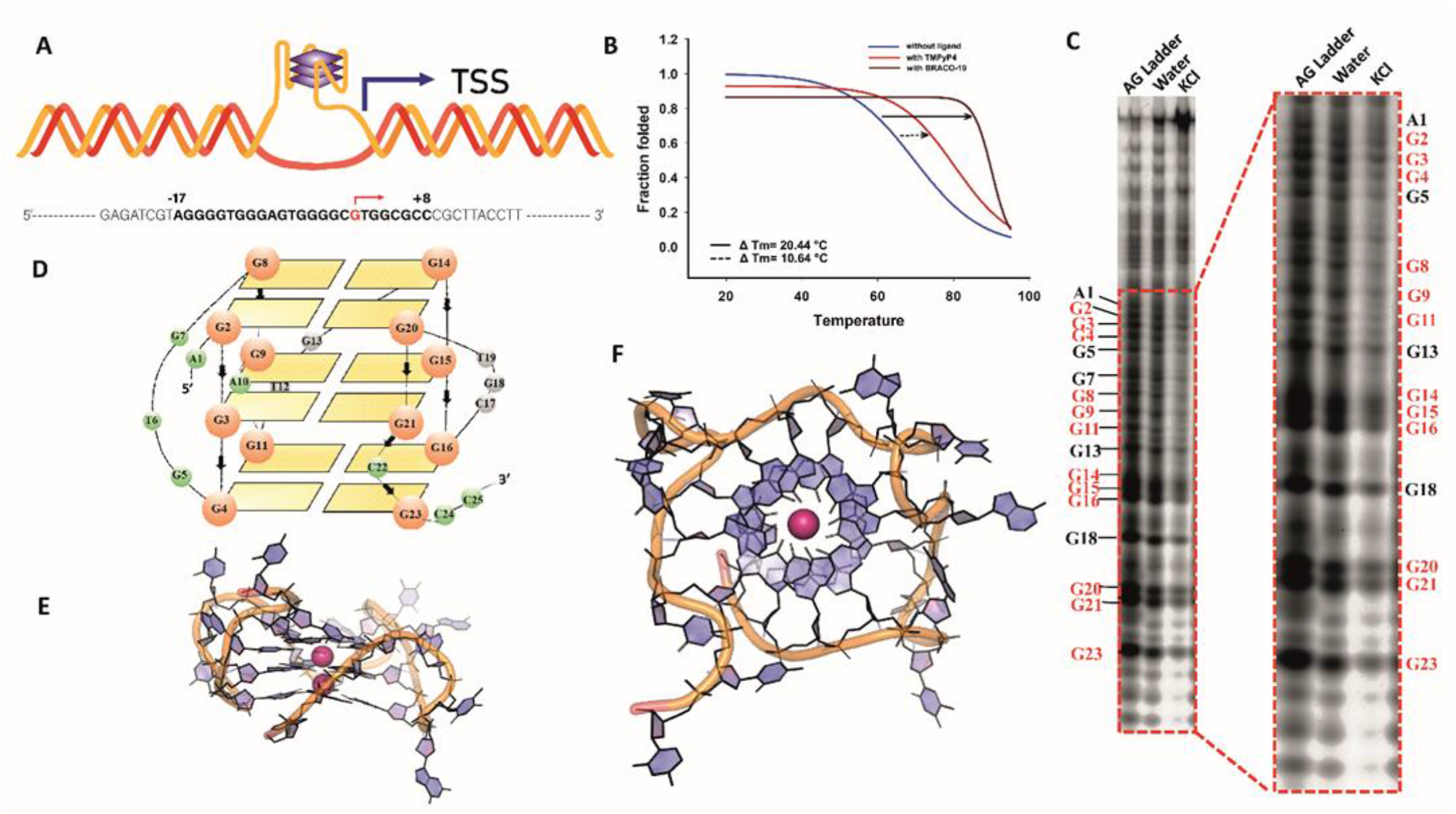
Biophysical characterization of SMARCAL1 P3 G4. (A) Schematic representation of SMARCAL1 promoter region with G4. (B) Change in the melting temperature of P3 G4 in the presence and absence of TMPyP4 and BRACO-19. (C) DMS foot-printing of P3 G4. (D) Schematic representation of the P3 G4. (E) Side view and (F) Top view of the P3 G4 model after energy minimization.

DMS footprinting of the G4 containing oligonucleotide revealed the specific guanine residues that might be involved in G-quartet formation (Figure 1C). The signal obtained from the experiment revealed the guanine residues that produced marked changes in signal intensity in water and ion containing experimental conditions which were analyzed by densitometric plot using ImageJ software (Figure S12). G2-G8-G14-G20 formed the first quartet, G3-G9-G15-G21 constituting the second quartet followed by G4-G11-G16-G23, which formed the third quartet. This G4 structure was found to contain three loops and two bulges as revealed from the footprinting experiment (Loop 1: G5•T6•G7•; Bulge1: A10; Loop 2: T12•G13•; Loop3: C17•G18•T19•; Bulge 2: C22•) (Figure 1D). This information was utilized to construct the parallel G4 model in the Maestro interface of Schrödinger software by editing the *c-Myc* G4 structure (PDB ID: 1XAV) appropriately. Minimization was performed in AMBER18 along with heating and equilibration, which produced the energetically minimized structure (Figure 1E &1F), revealing the stabilized G-quartet core containing the two central potassium ions coordinating with the O6 of guanine residues.

We used the porphyrin derivative, TMPyP4 [5,10,15,20-Tetrakis (1-methyl-4-pyridinio) porphyrin tetra (p-toluene sulfonate)] and acridine derivative BRACO-19 [N,N’-(9-{[4-(dimethylamino) phenyl] amino} acridine-3,6-diyl) bis (3-pyrrolidin-1-ylpropanamide)], two widely studied, G4 binding probe, to investigate the dynamic stability of the G4 structures in vitro.^24 25 26^ Significant changes in the intensity of the CD spectra was observed by titrating TMPyP4 (Figure S3) and BRACO-19 with the putative P3 G4 sequence, as both TMPyP4 and BRACO-19 interacted with the P3 G4 sequence to give rise conformational changes. CD melting data suggested that TMPyP4 and BRACO-19 conferred an elevation of the melting temperature of P3 G4 with a ΔTm of 10.64^°^C and 20.44^°^C respectively, thus suggesting the binding of TMPyP4 and BRACO-19 led to the higher stability of the G4 (Figure 1B). Further, we performed fluorescence spectroscopy to investigate the change in the emission spectra due to the interaction of P3 G4 with TMPyP4. We observed ∼45% hypochromic shift along with ∼7 nm red shift with increasing concentration of G4 sequence till saturation (Figure S4), thus signifying strong complex formation between P3 G4 and TMPyP4. So, the overall interaction studies suggested that the putative P3 G4 motif of SMARCAL1 promoter sequence can form parallel G4 structure and both TMPyP4 and BRACO-19 showed noteworthy interaction with P3 G4 sequence. See Supporting Information (SI) for details.

It is already found that, SMARCAL1 is associated with the ALT telomeres to resolve replication stress, thus providing telomere stability.^10^ Induction of telomere-specific replication stress is already investigated as a potent avenue to destroy ALT cancer cells.^4 27^ Thus, we investigated the potential role of the identified P3 G4 motif behind the downregulated expression of SMARCAL1 in ALT positive, osteosarcoma cell lines U2OS and Saos-2.^1 27^ The role of P3 G4 in the transcriptional regulation of SMARCAL1 was investigated by using TMPyP4 and BRACO-19. First, the transcriptional regulation was investigated at the mRNA level of SMARCAL1 by performing PCR assay upon treatment of 20, 40 and 60 μM TMPyP4 and 15 μM BRACO-19 for 24 h in U2OS and Saos-2 cell lines.^28 29 30^ 20 μM of TMPyP4 reduced SMARCAL1 expression by ∼ 60% in U2OS (Figure 2A). The SMARCAL1 mRNA expression was further reduced at 40 μM and 60 μM in both the cell lines (Figure 2A and S16).. Significant repression of ∼ 65% was also observed for 15 μM BRACO-19 treatment in U2OS cells (Figure 2B). In comparison, this transcriptional reduction was not observed in case of *ACTB or 18s rRNA* for not having any putative G4 forming sequence in its promoter. To confirm the *in cellulo* formation and stabilization of the *SMARCAL1* P3 G4 upon TMPyP4 treatment, we performed BG4 ChIP and checked for *SMARCAL1* G4 forming region. BG4 antibody binds to the G4 structure with high specificity.^31^

**Figure 2.**
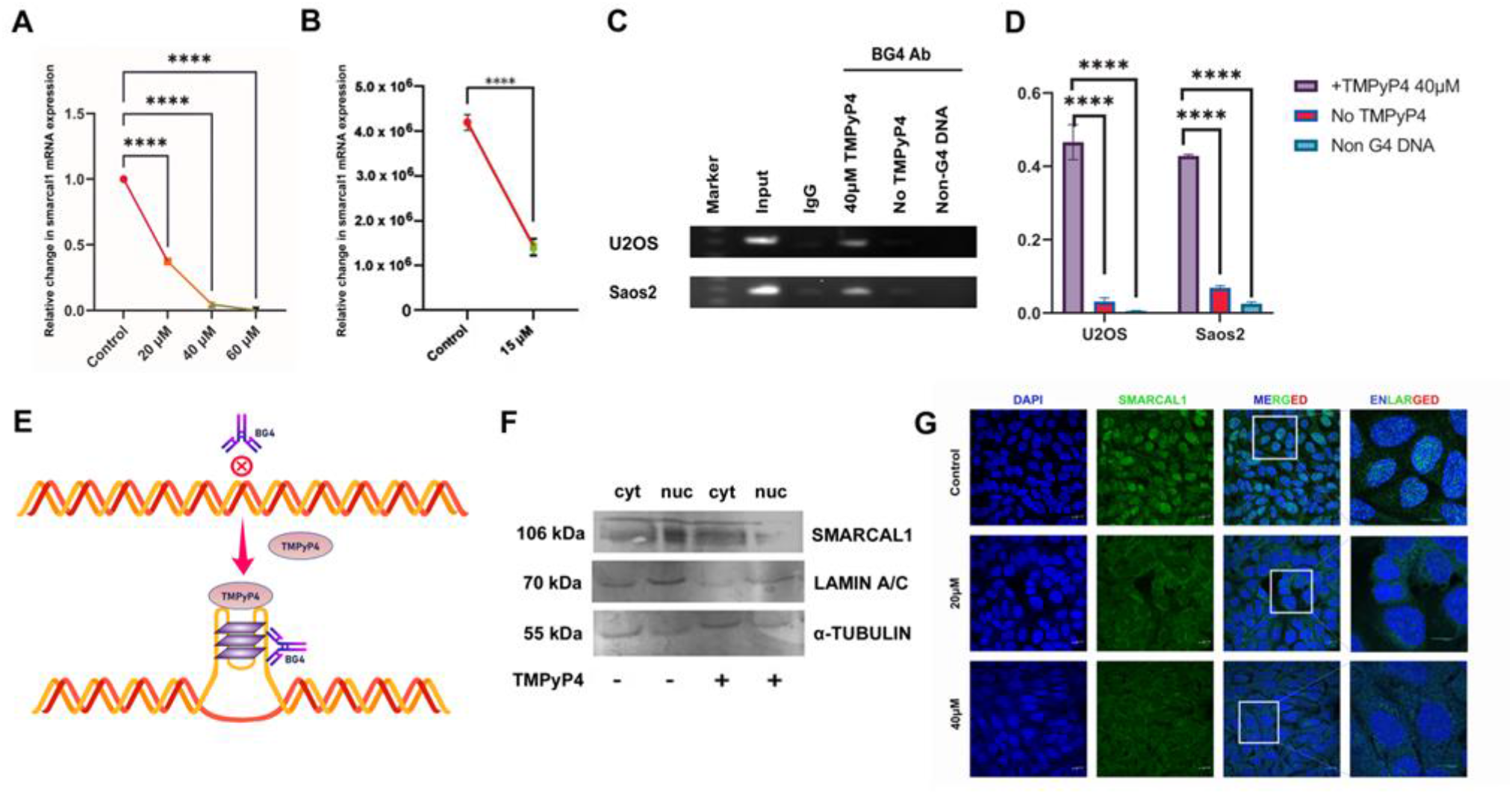
In cellulo expression profile of SMARCAL1 through ligand mediated G4 stabilization. SMARCAL1 mRNA expression was checked by PCR in U2OS cells, treated with TMPyP4 (A) and BRACO-19 (B) for 24 h. (C) BG4 ChIP experiment was performed in U2OS and Saos2 treated with 40 μM TMPyP4 targeting the G4 forming region in the promoter. (D) Graphical representation and (E) Schematic representation of BG4 ChIP, dictating G4 formation and stabilization in presence of TMPyP4. (F) Western blot dictating the change in the expression of SMARCAL1 protein in the cytosolic (cyt) and nuclear (nuc) regions of control, 40μM TMPyP4-treated U2OS cells. α-tubulin was used as loading control. (G) Immunostaining with antibody against SMARCAL1 (green) was done in U2OS cells, treated with 20 and 40 μM TMPyP4 for 24 h and compared with that of control. The nuclei stained with DAPI (blue) and merged images of SMARCAL1 and DAPI were also shown. Scale bar: 10 μm.

We observed significant G4 formation and stabilization in the presence of TMPyP4 in U2OS and Saos-2 cell lines, which was significantly reduced in the absence of TMPyP4 (Figure 2C and 2D) thus clearly suggesting that the G4 is being formed and stabilized by TMPyP4 *in cellulo*, thus interacting with BG4 antibody. No such stabilization and BG4 interaction was observed in the case of non-G4 DNA, which do not have any G4 motifs. We also noticed reduced expression at the protein level of SMARCAL1 in U2OS by western blot (Figure 2F) and confocal imaging (Figure 2G and Figure S7) of SMARCAL1 in nuclear region, which again proved the effectiveness of G4 ligands, i.e., TMPyP4 and BRACO-19 at this concentration in both the cell lines (see SI for details).

Later on, we continued to investigate the consequences of this SMARCAL1 downregulation in ALT positive cell line through different experimental sets by confocal microscopy. It was already reported that SMARCAL1 accumulates in ALT telomeres to resolve replication stress and provide telomere stability. Consistent with this, we also found that the localization of SMARCAL1 was drastically reduced from ALT telomeres in both the cell lines, with the increasing concentration of TMPyP4 (Figure S10 A & S10B). This SMARCAL1 downregulation can increase replicative stress at both telomere and genome level in ALT positive cells. Thus, we predicted that the downregulation of SMARCAL1 in ALT positive cells would induce accumulation of collapsed replication forks and increased PML localization at telomeres.^10 32^ By treating U2OS cells with the effective concentration of both TMPyP4 and BRACO-19, we observed increased localization of PML at ALT telomeres (Figure 3A, S8A, S8B and S14), thus suggesting the induction of replication stress at ALT telomeres due to the downregulation of SMARCAL1 through P3 G4 stabilization by TMPyP4 and BRACO-19, thus confirming the higher sensitivity of ALT positive cells to TMPyP4 and BRACO-19-induced replication stress in ALT telomere regions. Increased PML localization at ALT telomeres was also observed in case of siRNA mediated silencing of SMARCAL1 gene, thus highlighting the efficiency of G4 ligands to stabilize the SMARCAL1 P3 G4 and further downregulate the SMARCAL1 expression (Figure 3A and S14). We also found a significant increase in DNA damage at genomic level due to the downregulation of SMARCAL1 in U2OS. (Figure S11A, S11B and S11C) See SI for details.

**Figure 3.**
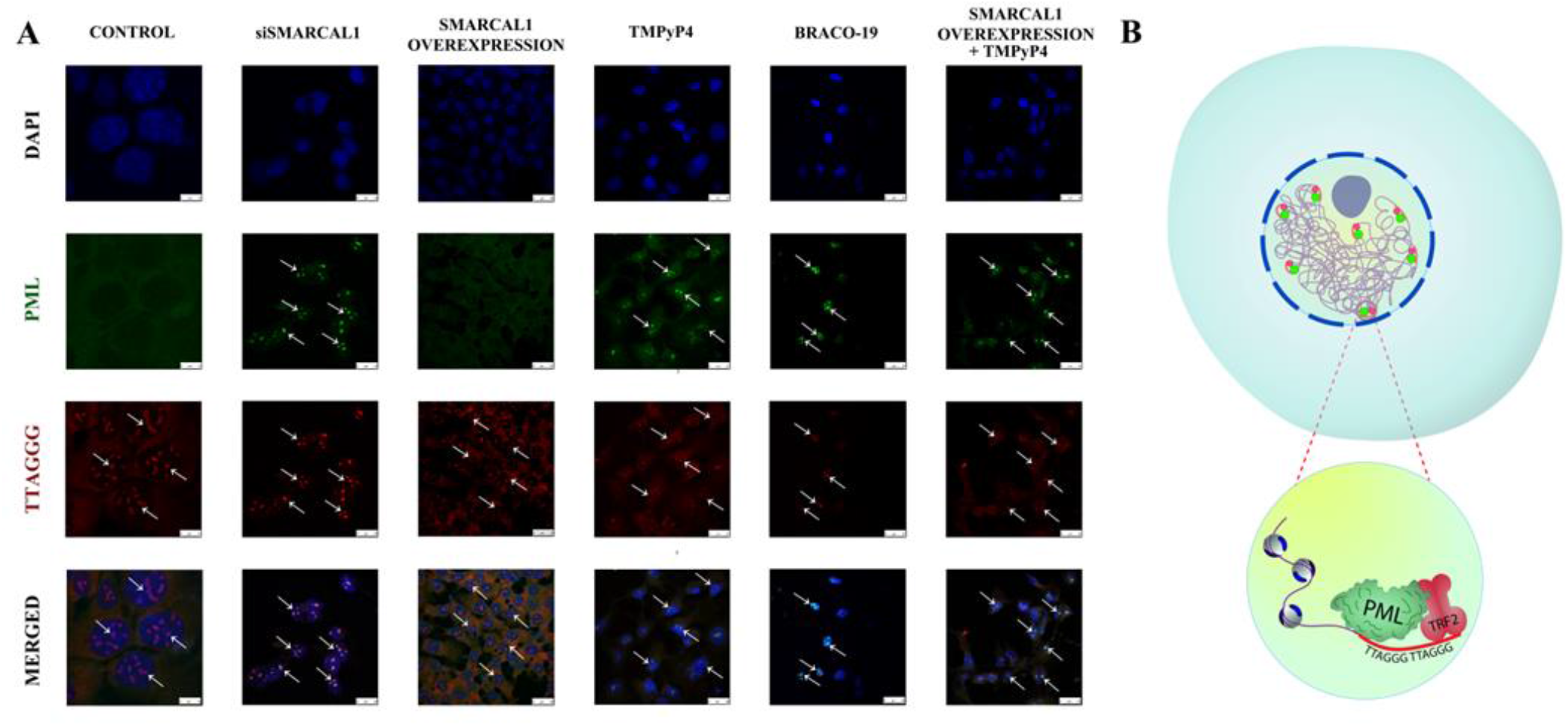
Induction of replication stress at ALT telomeres by SMARCAL1 downregulation. (A) Cells were treated with 40 μM TMPyP4 and 15 μM BRACO-19 for 24 h and compared with that of control, siSMARCAL1 and SMARCAL1 overexpressed conditions. Combined immunofluorescence and fluorescence in situ hybridization (FISH) analyses the co-localization of PML at telomeres (TTAGGG) in U2OS cells. The nuclei stained with DAPI (blue) and merged images of PML (green) and telomere (red) were shown. Representative images were shown. (B) Schematic representation of PML localization at ALT telomere regions.

In addition, as TRF2 specifically interacts with the TTAGGG duplex repeats at human telomeres and protects telomeres from end-to-end fusions by the formation of the t-loop, we also investigated the probable interaction between PML bodies and TRF2 at telomeric repeats due to SMARCAL1 downregulation.27 28 29 Consistent with the PML and TTAGGG co-localization data, here also we found that PML bodies are interacting with TRF2 at telomere regions.^2 33 34^ Both U2OS and Saos-2 displayed a significant rise in TRF2 and PML co-localization in the telomere region within nuclear periphery upon increasing concentration of TMPyP4 (Figure S9A and S9B). No such significant rise in co-localization was observed in untreated ALT cells, supporting pivotal role of SMARCAL1 promoter G4 and its stabilization through TMPyP4, which again confirmed that, G4 ligands mediated downregulation of SMARCAL1 induces increased accumulation of APBs in ALT telomere repeat regions recruiting TRF2, thus indicating replication stress, telomere instability and telomere dysfunction in ALT positive cells.

In the last few decades, different G4 ligands have been reported to show anti-cancer activity in telomerase dependent cancers and also in ALT-positives.^35 36 37^ Recent research already reported the possibility that G4 stabilization can inhibit the ALT-based recombination pathway through the hyperactivation of

ALT.^38 39^ Downregulation of another ATP dependent translocase FANCM resulted in the activation of hyper-recombinogenic phenotype in ALT, thus inducing apoptosis in ALT positive cancers.^4 40 41^ Interestingly, G4 stabilization has already been reported as a reliable strategy for the selective targeting of ATRX-deficient ALT positive gliomas.^42^ In this work, we found that SMARCAL1 downregulation led towards the co-localization of PML at ALT telomere regions and development of APBs, a potent marker of ALT, thus reflecting higher replication stress and hyper-recombinogenic phenotype in ALT cells. It is already suggested that the hyper-activation of ALT along with high levels of DNA damage and APBs may induce apoptosis. Recently, it has been reported that SMARCA4 (BRG1), a SMARCA-type protein binds at G4 structures in chromatin.^43^ Moreover, BRG1 and SMARCAL1 can mutually co-regulate each other to increase their transcript levels during DNA damage.^44^ In this situation, BRG1 binds to SMARCAL1 promoter, leading towards an increase in SMARCAL1 transcripts. However, as a result of the G4 ligands mediated stabilization of SMARCAL1 G4, we observed a significant downregulation of the SMARCAL1 transcripts, suggesting that BRG1 occupancy on the SMARCAL1 promoter may not be sufficient for SMARCAL1 expression and thus emphasizing the crucial regulatory function of the SMARCAL1 promoter G4 motif. Therefore, induction of hyper-recombinogenic phenotype in ALT through SMARCAL1 downregulation by TMPyP4 and BRACO-19 mediated G4 stabilization may selectively destroy ALT cancers by introducing telomere replication stress. Further investigations are needed to comment on the type of ALT pathway that is being activated due to the SMARCAL1 downregulation.

Sustained telomeric replicative stress is required to maintain ALT, which is administered by SMARCAL1. When excessive, it can trigger apoptosis in ALT cells. Fueling of ALT mediated by G4 stabilizing molecules already highlighted their possible application in a new therapeutic window to target ALT-positive tumors.^27 38 39 45^ So, in accordance with this, our study will surely put SMARCAL1 as a new, potential target in the map of future ALT therapeutics. Moreover, this study will also open a new avenue to target ALT-associated G4 structures for the development of selective G4 based ALT-therapeutics.

## ASSOCIATED CONTENT

### Supporting Information

The Supporting Information is available free of charge at

Detailed description of materials and methods, supplementary tables, supporting discussions, other supplementary Figures with figure legends and additional references. (PDF)

## AUTHOR INFORMATION

### Authors

**Suman Panda** - Department of Biophysics, Bose Institute, EN 80, Sector V, Bidhan Nagar, Kolkata – 700091, WB, India.

**Tanaya Roychowdhury**- Sloan Kettering Institute, Memorial Sloan Kettering Cancer Center, 1275 York Ave, New York, NY 10065, United States of America.

**Anindya Dutta**- Department of Biophysics, Bose Institute, EN 80, Sector V, Bidhan Nagar, Kolkata – 700091, WB, India.

**Sourio Chakraborty**- Division of Molecular Medicine, Centenary Campus, Bose Institute, P-1/12 C.I.T. Scheme VII-M, Kolkata - 700054, WB, India

**Tanya Das**- Division of Molecular Medicine, Centenary Campus, Bose Institute, P-1/12 C.I.T. Scheme VII-M, Kolkata - 700054, WB, India.

## Author Contributions

SP: Formal analysis, Investigation, Methodology, Software, Visualization, Writing – original draft, Writing - review & editing. TR: Formal analysis, Investigation, Methodology, Software, Visualization. AD: Formal analysis, Methodology, Writing - review & editing, Software and Visualization. SC: Formal analysis, Investigation, Software, Visualization. TD: Funding, Supervision, Validation, Resources and Project administration. SC: Conceptualization, Funding acquisition, Formal analysis, Supervision, Validation, Resources, Project administration, Writing - review & editing. All authors have given approval to the final version of the manuscript. ‡These authors contributed equally.

## ACKNOWLEDGMENT

The instrumental support received from the laboratory of S. Sau; Bose Institute is highly acknowledged. We are thankful to the cell culture facility of the laboratory of S. Chattopadhyay, CSIR-IICB and the computational facility of the laboratory of A. Bhunia, Bose Institute. Central instrument facilities of Bose Institute and CSIR-IICB are also highly acknowledged.

## ABBREVIATIONS

ACTB: beta-actin
CD: Circular dichroism
TRF2: Telomeric repeat-binding factor 2
ChIP: Chromatin immunoprecipitation
ATRX: alpha-thalassemia/mental retardation, X-linked.

---

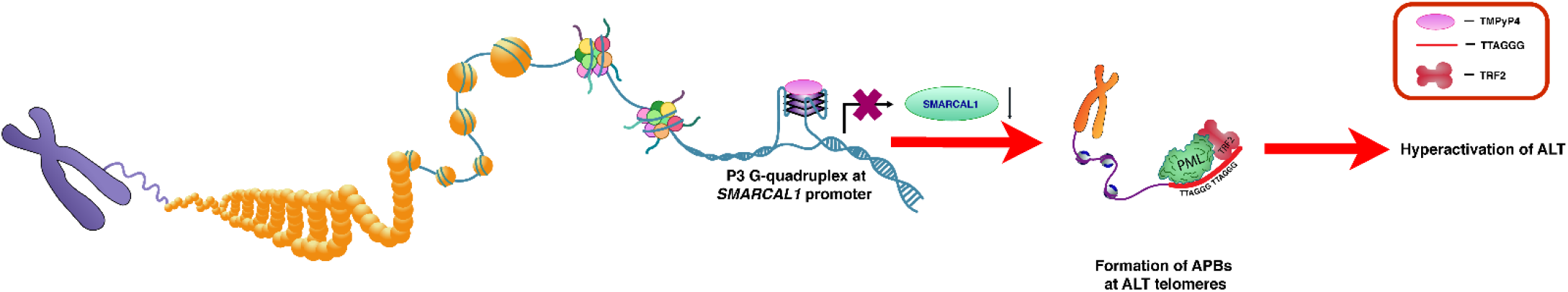

