## Supporting Information for "ALTering cancer by triggering telomere replication stress through the stabilization of promoter G-quadruplex in *SMARCAL1*"

**ABSTRACT:** Most of the human cancers are dependent on telomerase to extend the telomeres. But ~10% of all cancers use a telomerase-independent, homologous recombination mediated pathway called alternative lengthening of telomeres (ALT). Due to poor prognosis, the ALT status is not being considered yet in the diagnosis of cancer. No such specific treatment is available till date for ALT cancers. ALT positive cancers are dependent on replication stress to deploy DNA repair pathways to the telomeres to execute homology recombination mediated telomere extension. SMARCAL1 (SWI/SNF related, matrix-associated, actin-dependent regulator of chromatin, subfamily A-like 1) is associated with the ALT telomeres to resolve replication stress thus providing telomere stability. Thus, the dependency on replication stress regulatory factors like SMARCAL1 made it a suitable therapeutic target for the treatment of ALT positive cancers. In this study, we found a significant downregulation of SMARCAL1 expression by stabilizing the G-quadruplex (G4) motif found in the promoter of *SMARCAL1* by potent G4 stabilizers, like TMPyP4 and BRACO-19. SMARCAL1 downregulation led towards the increased localization of PML (promyelocytic leukaemia) bodies in ALT telomeres and triggered the formation of APBs (ALT-associated promyelocytic leukaemia bodies) in ALT positive cell lines, thus increasing telomere replication stress and DNA damage at genomic level. Induction of replication stress and hyper-recombinogenic phenotype in ALT cells mediated by G4 stabilizing molecules already highlighted their possible application as a new therapeutic window to target ALT-positive tumors. In accordance with this, our study will also provide a valuable insight towards the development of G4 based ALT-therapeutics targeting *SMARCAL1*.

#### Table of contents

**1. Material & Methods** **S3-S11**

**2. Supplementary tables (S1 to S8)** **S12-S13**

**3. Supporting discussions** **S14-S15**

**3.1. Thermodynamic profiles of TMPyP4 and BRACO-19 with P3 G4.**

**3.2. *In cellulo* effect of TMPyP4 on the promoter G-quadruplex of *SMARCAL1***

**3.3. Downregulation of SMARCAL1 induces DNA damage throughout the genome.**

**4. Supplementary Figures and Figure legends** **S16-S40**

**Figure S1.** CD Spectra of the wild type oligonucleotide sequence (P3 wt).

**Figure S2.** CD Spectra of the mutated oligonucleotide sequence (P3 mut).

**Figure S3.** CD spectra of P3 wt due to TMPyP4 interaction.

**Figure S4.** Fluorescence emission spectra of TMPyP4 titrated with increasing concentration of G4 forming sequence (P3 wt) till saturation.

**Figure S5.** ITC profiles of P3 wt with TMPyP4 and BRACO-19.

**Figure S6.** Dual-luciferase assay.

**Figure S7.** Immunocytochemistry of SMARCAL1.

**Figure S8.** Recruitment of PML bodies (APB foci) at ALT telomeres due to SMARCAL1 downregulation.

**Figure S9.** Interaction profile and co-localization of PML and TRF2 protein.

**Figure S10.** Loss of SMARCAL1 accumulation in ALT telomeres due to TMPyP4 mediated stabilization of the SMARCAL1 G-quadruplex.

**Figure S11.** Loss of SMARCAL1 induces DNA damage at genomic level.

**Figure S12.** DMS footprinting densitometric plot.

**Figure S13.** MTT assay for BRACO-19 in U2OS cells.

**Figure S14.** Graphical representation of the APBs formation in different experimental sets.

**Figure S15.** Expression profile of SMARCAL1 mRNA in U2OS cells.

**Figure S16.** Graphical analysis of SMARCAL1 mRNA expression profile in Saos-2 cells.

**Figure S17.** Graphical representation of the fluorescence intensity of SMARCAL1 protein.

**Figure S18.** Densitometric analysis of the Western Blot.

**5. Additional references** **S41**

### 1. Materials and methods

#### 1.1. Bioinformatics Analysis

The complete *SMARCAL1* gene sequence was obtained from NCBI/Ensembl (Gene ID: 50485; Ensembl:ENSG00000138375; MIM:606622). Complete characterization of the gene sequence was done to obtain the coding regions. The promoter region of the gene was predicted by using MatInspector from the Genomatrix software suite. The putative G-quadruplex motif was analysed by QGRS mapper (http://bioi nformatics.ramapo.edu/QGRS/index.php; Supplementary Table S1). The putative transcriptional start site (TSS) of *SMARCAL1* gene was determined from Eukaryotic Promoter Database (EPD) of the Swiss Institute of Bioinformatics (SIB) (https://epd.epfl.ch//index.php).

#### 1.2. Oligonucleotide sequences and chemicals

The oligonucleotide sequences for the biophysical experiments were purchased from Eurofins India Pvt. Ltd. (Supplementary Table S2) and dissolved in water. The cationic porphyrin derivative, 5,10,15,20-Tetra-kis (1- methyl-4- pyridinio) porphyrin tetra (p-toluenesulfonate; TMPyP4) and acridine derivative BRACO-19 [N,N'-(9-{[4-(dimethylamino)phenyl]amino}acridine-3,6-diyl)bis(3-pyrrolidin-1-ylpropanamide)], were obtained from Sigma-Aldrich (catalog no. 323497 and SML0560 respectively) and dissolved into ultrapure water (Invitrogen) to make a stock concentration of 10 mM and stored in dark at 20 ◦C. For *in vitro* biophysical interaction studies, TMPyP4 was diluted to different working concentrations and allowed to equilibrate at room temperature for an hour. All the oligonucleotide sequences were annealed at 95 ◦C for 5 min at specific KCl concentrations (10 mM potassium phosphate buffer mixed with required amount of KCl) as required and allowed to cool down slowly to room temperature and stored in aliquots at 20 ◦C.

##### 1.3. Circular Dichroism Spectroscopy

CD spectra of all the putative G4 oligonucleotide sequences were being recorded using a Jasco-815 Circular Dichroism spectrophotometer at 25 ◦C in the wavelength range of 300–210 nm with a scan speed of 100 nm/min, step size of 1 nm and a bandwidth of 1 nm. All the measurements were carried out using a standard quartz cuvette having path length of 1 mm and total reaction volume of 350 μl. The CD melting experiments were performed using the same cuvette and buffer in the temperature range of 20–95 ◦C at an interval of 5 ◦C. The samples were incubated for three minutes before all the measurements. The experiments were carried out using 10 mM potassium phosphate buffer containing 100 mM potassium chloride at pH 7. All the experiments were carried out using oligonucleotide concentration of 20 μM and were also titrated using increasing concentration of TMPyP4 and BRACO-19 up to a maximum ratio of 1:3 till the saturation reached. All the readings were noted after 5 min of incubation to ensure stable complex formation after binding. The experiments were performed twice. The plot of fraction folded vs temperature was fit with the sigmoidal three parameter equation for P3 G4 sequence using the equation

$$y = \frac{a}{1 + e^{-\left(x - x0/_{b}\right)}}$$

The equation for Gaussian three parameters was used for the fitting of P3 CD melting curve in presence of TMPyP4. The equation follows:

$$y = ae^{\left[-0.5\left(x - x0/_{b}\right)^{2}\right]}$$

##### 1.4. Fluorescence Spectroscopy

Fluorescence measurements were carried out using Jasco FP 8200 spectrofluorometer (Tokyo, Japan) to observe the binding profile of TMPyP4 with the putative G4 sequence of *SMARCAL1*. Fluorometric titrations were performed in a 10 mm quartz cell at 25ºC. The emission spectra at each point of titration were collected between 600 – 750 nm having both excitation and emission slits defined at 5 nm. Fluorescence titrations were carried out by stepwise addition of oligonucleotide sequences to the cell containing 10 μM ligand in potassium phosphate buffer (pH 7.0) till saturation. Five minutes of incubation time was provided after each titration for the equilibration of the complex. Excitation wavelength (λex) was 435 nm for TMPyP4.

##### 1.5. Isothermal Titration Calorimetry

To unravel the binding thermodynamics, ITC experiments were performed in an Affinity ITC, TA instruments, at 25°C temperature. All the samples were prepared through extensive degassing before the experiment. TMPyP4 (400 μM) and BRACO-19 (500 μM) were taken in the syringe and were injected into the fixed concentration of oligonucleotide solution (10 μM) taken in the cell in two different experiments. After deducting the blank, dissociation constants were determined by titrating TMPyP4 and BRACO-19 with 10 μM of P3 G4 motif containing sequence. The binding isotherm was analyzed using the NanoAnalyze v3.10.0 software from TA instruments. After fitting the isotherm, equilibrium association constant ($K_a$), enthalpy (ΔH) and entropy (ΔS) of the binding were obtained. $\Delta G$ of the reaction was calculated from the thermodynamic parameters using the following equation:

$$\Delta G = \Delta H - T\Delta S$$

##### 1.6. Dimethylsulfate (DMS) protection assay

6-FAM labelled oligonucleotides at the 3′ termini including the putative G4 stretch was purchased from Sigma-Aldrich (Supplementary Table S3) and dissolved in TE buffer (pH = 8) at 100 μM concentration to prevent hydrolysis. The annealing of 6-FAM labelled oligonucleotide was done using a sub-stock concentration of 10 μM at 95 ◦C for 5 min in both potassium phosphate buffer (10 mM; pH = 7.0) mixed with 100 mM KCl and in ultrapure water (used as negative control) separately. Both were gradually cooled to room temperature and kept overnight. DMS reaction of the annealed oligonucleotides was carried out using 1% DMS (Merck) for 1 min in 100 μl reaction mixture. Then, 25 μl of freshly prepared stop solution (50 μl sterile water, 43 μl sodium acetate (pH = 7.0), 7 μl β-mercaptoethanol) was added immediately to the reaction mixture to stop the reaction. Next, 250 µl chilled ethanol and freshly prepared sodium acetate (pH = 5.2, 0.3 M) was added immediately after stopping the reaction. After that, the sample was centrifuged at 13,000 rpm for 30 min at 4 ◦C. The pellet was washed with 1 ml of freshly prepared 70% ethanol and then lyophilized in a Biobase lyophilizer instrument. The samples were then treated with 1 M piperidine (Sigma) and incubated at 95 ◦C for 20 min. After that, the samples were washed thrice with sterile water and lyophilized each time to remove all the traces of piperidine from the samples. In a separate reaction, the A+G ladder was also prepared by incubating 5 µl of sample dissolved in water with 4% formic acid (Merck) for 30 min at 37 ◦C and lyophilized immediately. The cleaved products after DMS reaction and A+G ladder were separated by running them in 15% denaturing PAGE sequencing gel. For analysis, gel images were taken in Amersham Typhoon 5 Biomolecular Imager.

##### 1.7. Modelling and Molecular Dynamics Simulation of G-quadruplex structure

Modelling of the *SMARCAL1* G4 structure was performed using *cMYC* promoter G4 NMR structure as the template (PDB ID: 1XAV). Residues were edited in Maestro GUI of Schrodinger Molecular Modelling Suite. The atom names and atom types were renamed in PDB editor. This model was taken up for minimization in AMBER 14 using the ff14SB and parmBSC1 forcefields for DNA. $K^+$ ions were placed to neutralize the negative charge of guanine residues. The system was incorporated in explicit solvent using TIP3P water box with 14.0 Å dimension on all the three axes (x, y and z-axis), to mimic solvent environment conditions inside cellular systems.

##### 1.8. Cell culture and treatment

ALT positive cancer cell lines, U2OS (ATCC HTB-96) and Saos-2 (ATCC HTB-85) and were purchased from American Type Culture Collection (ATCC). U2OS cell line was grown and maintained in McCoy's 5 medium modified (Thermo Fisher), supplemented with 10% (v/v) fetal bovine serum (Gibco), 100 U/ml penicillin (Gibco), and 100 µg/ml streptomycin (Gibco) at 37◦C in 5% (*v*/v) $CO_2$. Saos2 cell line was grown and maintained in McCoy's 5A medium modified (Thermo Fisher), supplemented with 15% (v/v) fetal bovine serum (Gibco), and 1% penicillin/streptomycin (Gibco) at 37◦C in 5% (*v*/v) $CO_2$. The cell lines were treated with

TMPyP4 and BRACO-19 at specific concentration for all the experiments as mentioned in results.

##### 1.9. Reporter constructs for luciferase vectors and over-expression clone

The promoter region of human *SMARCAL1* gene with putative G4 forming sequence was amplified by PCR using human genomic DNA as template and then cloned into pGL4.72[*hRlucCP*] vector (Promega; Madison, USA; catalog no. E6901) at *Kpn*I/*Hin*dIII site. This construct was named as pSMARCAL1-WT, contained 332 bp (-239 to +93 from TSS) sequence with the putative G4 forming stretch of *SMARCAL1*. Its mutation variant, named as pSMARCAL1-Null, was synthesised by deleting the G4 forming stretch only. Mutated construct was outsourced from Dr. KPC Life Sciences Pvt. Ltd., India. For SMARCAL1 over-expression studies, SMARCAL1 (NM_001127207) Human Tagged ORF Clone (Cat. no.: RC226275) was procured from OriGene, USA.

##### 1.10. Luciferase assay

For luciferase reporter assay, approximately $4 \times 10^3$ cells were seeded on 96-well culture plate. After 24 hours, each pGL4.72[*hRlucCP*] construct (wild-type and mutated; 100 ng) was transfected transiently into cells with Lipofectamine 2000 (Invitrogen). After 4 hours of transfection, the medium was replaced with fresh medium with appropriate doses of TMPyP4 (for treated cells) or PBS (for control cells). After that, the cells were further incubated for next 24 hours. Each transfection was normalized with pGL3 vector (10 ng; Promega). Then the cells were harvested and lysed with 1× Passive Lysis buffer (Promega) to get the Firefly and *Renilla* luciferase activities using Dual-Luciferase Reporter Assay System (Promega) in a luminometer. For each sample, *Renilla* luciferase activity was normalized with Firefly luciferase activity and their ratio was considered as relative luciferase activity. After further normalization of all the TMPyP4 treated samples with control, the fold change of mutant is represented. The fold change of wild type is calculated with respect to the mutant values.

##### 1.11. Quantitative Real-time PCR (q-PCR)

Following the standardized protocol, total RNA was isolated from cells using RiboZol™ RNA extraction reagent (Amresco). cDNA synthesis was done using RevertAid fast strand cDNA synthesis kit (Thermo Scientific) and then q-PCR assay was performed by using Power SYBR GREEN mix (Invitrogen). The comparative $C_T$ method ($\Delta\Delta C_T$) was used to measure relative gene expression and the fold enrichment was calculated as: $1/2^{\Delta\Delta CT}$. Here, $\Delta C_T$ is the $C_T$ of target gene subtracted from the $C_T$ of the housekeeping gene, actin. $\Delta\Delta C_T$ is the $\Delta C_T$ of control sample subtracted from $\Delta C_T$ of treated sample. The primer sequences used in this qPCR are mentioned in Supplementary Table S4.

**1.12. Nuclear Cytosolic fractionation and western blotting**

Control and TMPyP4-treated U2OS cells grown in 10-cm tissue culture dishes were harvested at 70% confluency. After removing culture medium from the dish, cells were washed twice with 5 ml of ice-cold phosphate buffer saline (PBS) pH 7.4, scraped from culture dishes using a cell lifter, and collected into a centrifuge tube. After brief centrifugation (13,500 g, 4°C, 10 s), supernatant was removed and cell pellet was resuspended in 900 μl of the Nuclear/Cytosolic extraction Buffer A (pH=7.9) (10mM KCl, 10mM HEPES-NaOH, pH=7.9, 0.1 mM EDTA, 1mM DTT, 1X protease inhibitor cocktail), briefly vortexed, incubated on ice for 15 mins, treated with 10% (v/v) NP-40 solution, followed by centrifugation at 13,000 rpm for 5 mins at 4°C. Supernatant was collected as the "cytosolic fraction" and 250 μl of Nuclear/Cytosolic extraction Buffer B (0.4 M NaCl, 20mM HEPES-NaOH, pH=7.9, 10mM EDTA, 1mM DTT, 1X protease inhibitor cocktail) was added to the cell pellet and resuspended well. After incubation on ice for 30 mins centrifugation was done at 13,000 rpm for 5 mins at 4°C. Supernatant was collected as the "nuclear fraction". After measuring protein concentration, 120 μg of total protein was loaded and electrophoresed on 8% SDS-PAGE and transferred on to PVDF membrane followed by blocking with non-fat skimmed milk (5%) for 1 hour and after that probed with specific primary antibody of SMARCAL1 (D3P5I, Cell Signaling Technology), LAMIN A/C (nuclear marker) (Cell Signaling Technology), and alpha-TUBULIN (cytosolic marker) (Thermo Scientific) for overnight at 4°C. After incubating with AP-conjugated secondary antibody, blots were developed using Alkaline Phosphatase method (HiMedia, India) and images were captured using GelDoc XR+ system.

##### 1.13. Confocal microscopy

Cells were grown ($1 \times 10^5$ cells/well) on glass coverslips in 6 well plates (BD falcon) using and grown upto 70% confluency. Then, cells were treated with appropriate concentration of TMPyP4 and BRACO-19 for 24 hours. Briefly, the media was discarded, and the coverslips washed thrice with 1× PBS (pH 7.4). After that, the cells were fixed with 4% paraformaldehyde for 15 mins, and again washed thrice with 1× PBS, followed by permeabilization with 1% Triton X-100 (PBST) for 10 min and blocked with 3% BSA for 1 hour. Cells were then incubated with anti-SMARCAL1 (CST), anti-γ-H2AX (CST), anti-TRF2 (Abcam) and anti-PML (Abcam) antibodies for 1 hour in different experimental sets. After washing with 1×PBS-T buffer for 5 mins for three times, the cells were incubated with secondary antibody for 1 hour at room temperature. Then the coverslips were washed with PBST and mounted with Prolong Gold Antifade with DAPI (Invitrogen). The images were captured using Leica TCA SP8 confocal microscope (Leica S5 Microsystems, Wetzlar, Germany) with Leica Application Suite (LAS). Images shown were the representatives of three independently performed experiments.

##### 1.14. Immuno-FISH (Fluorescence In-Situ Hybridization)

U2OS and Saos2 cells were grown on glass cover slips at a density of 0.6 million cells in complete DMEM medium in a 37 °C incubator. Next day cells were washed with 1x PBS thrice. Cells were fixed with 4% paraformaldehyde (PFA) for 10min at RT. PFA fixed cells were incubated with 0.2% triton X 100 in PBS for 15min and then washed thrice with PBS (5min, each). Blocking was done using 1xPBG (PBS, BSA, Gelatin) for 1h at RT. The primary antibody was diluted in 1X PBG (1: 250), added to coverslip and incubated overnight at 4°C. Next day coverslips were washed with 1xPBS thrice and incubated with secondary antibody (1:1000) for 2h at RT. Post incubation coverslips were washed with 1xPBS thrice. Cells were re-fixed with

4% PFA+ 0.1% triton X 100 for 15min at RT followed by incubation in 10mM glycine in $H_2O$ for 30min at RT. Coverslips were washed with 1XPBS thrice (5min each). Cells were treated with RNase A 200mg/ml for 30min followed by dehydration in a series of ethanol washes 70%, 85%, and 100%. The coverslips were then dried. 10nM Telomeric FISH probe in hybridization buffer (50% formamide, 2XSSC, 2mg/ml BSA, 10% dextran sulfate) was added to coverslips and DNA was and then placed in a humidified chamber overnight. The coverslips were then washed in 2X SSC +50% formamide, 2X SSC alone and finally in 2X SSC containing DAPI. Coverslips were mounted on glass slides with Prolong Gold antifade and allowed to cure overnight. Imaging was done using Leica SP8 confocal microscope.

##### 1.15. BG4 Chromatin immunoprecipitation (ChIP)

U2OS and Saos2 cells were seeded in 10cm dishes in DMEM complete medium with 10% Fetal Bovine Serum (GIBCO) and 1% antibiotic and anti-mycotic and incubated at 37°C at 5% $CO_2$. Cells were either mock treated or treated with 40µM TMPyP4 for 24h. Next day cells were fixed using 2% (v/v) formaldehyde for 10 min at RT. Quenching was done using 125 mM glycine for 5min, cells were pelleted and washed twice with ice cold PBS containing 1X PIC and PI. The flash-frozen pellets were lysed for 10 min on ice in 100 µl of 50 mM Tris-HCl pH 8.0, 10 mM EDTA, 0.5% SDS, and protease inhibitor cocktail. Samples were sonicated on ice using the optimized condition to shear chromatin to an average size of 100–500 bp (10 pulses of 30s each with intermittent 30s rest on ice). Sheared chromatin was diluted 1:5 in IP-buffer (10 mM Tris-HCl pH 7.5, 1 mM EDTA, 0.5 mM EGTA, 1% Triton X-100, 0.1% SDS, 0.1% Na-deoxycholate, and 140 mM NaCl) supplemented with protease inhibitor cocktail. After centrifuging 10 min at 15,000 rpm at 4 °C for 10min, the supernatant containing soluble chromatin fraction was recovered and incubated with 0.7 mg/ml RNase A (ThermoFisher) for 30 min at 37 °C. For chromatin immunoprecipitation, 10 µl protein-A/G beads (Pierce™ ThermoFisher) were washed in IP-buffer and incubated with 1 µg Anti-FLAG Ab (Sigma Aldrich #F3165) for 1 h at 4 °C on a rotating wheel. A total of 50 µl of RNA digested chromatin were incubated with BG4 Ab (Merck Millipore #MABE917 or without for the Mock negative control) overnight at 4°C. The anti-FLAG-coated beads were washed with IP-buffer and incubated with chromatin–BG4 complex for 3 h at 4 °C on a rotating wheel. Beads were washed four times with IP-buffer and once in wash buffer (10 mM Tris-HCl pH 8.0 and 10 mM EDTA). Elution of immune-precipitates and chromatin crosslink reversal were performed incubating beads with 70 µl elution buffer (10 mM Tris-HCl pH 8.0, 5 mM EDTA, 300 mM NaCl, and 0.5% SDS) containing 0.3 mg/ml RNase A (ThermoFisher) for 30 min at 37 °C followed by the addition of 0.5 mg/ml proteinase K for 1 h at 55 °C. Sample were then transferred to 100ºC in dry bath and incubated for 10min to inactivate the enzyme followed by centrifugation at 13,000 rpm for 10min at 4°C. Phenol chloroform extraction was performed followed by DNA precipitation using 3M sodium acetate, pH 5.2 Samples were stored at -20ºC overnight followed by pellet wash in 70% ethanol. The dried DNA pellet was resuspended in suitable volume of 1x TE buffer. Primers were designed against the G4 sequence of the *SMARCAL1* promoter region and a non-G4 forming region. Semi quantitative PCR was performed to analyze changes in response to TMPyP4 treatment. Experiments were done in triplicates.

**1.16. Statistical analysis**

Statistical analysis was performed using Microsoft excel. All the data for specific experiments were expressed as mean SD of three independent replicates. The SD was represented by error bars. The statistical significance between groups was calculated by two-tailed Student's *t*-test using GraphPad software (San Diego, USA). Paired t-test was calculated using Microsoft Excel. $p < 0.05$ was considered as significant. * p < 0.05, **p < 0.01, ***p < 0.001, ****p < 0.0001, ns, not significant.

#### 2. Supplementary tables

**Table S1:** The sequence of putative *SMARCAL1* G4 forming motif and QGRS Mapper analysis (with G-score):

| Sl. No. | Name | Sequence (5'-3') | Length | G-score |
|---|---|---|---|---|
| 1. | P3 | AGGGGTGGGAGTGGGGCGTGGCGCC | 25 | 21 |

**Table S2:** The sequence of oligonucleotides used in biophysical study:

| Sl. No. | Name | Sequence (5'-3') | Length |
|---|---|---|---|
| 1. | P3 Wild type | AGGGGTGGGAGTGGGGCGTGGCGCC | 25 |
| 2. | P3 Mutated | AGTTGTGTGAGTGTTGCGTGTCGCC | 25 |

**Table S3:** The sequence of oligonucleotide used in DMS footprinting study:

| Sl. No. | Name | Sequence | Length | Modifi-cation |
|---|---|---|---|---|
| 1. | P3 | GAAGTGGGCCAATGGGAAGGGTGAATCCAAGTGGAGATCGTAGGGGTG GGAGTGGGGCGTGGCGCCCGCT | 70 | 6-FAM at 3' |

**Table S4:** The sequences of the oligonucleotide primers used in q-PCR:

| Sl. No. | Gene Name | Forward primer (5'- 3') | Reverse primer (5'- 3') |
|---|---|---|---|
| 1. | ***SMARCAL1*** | TCCCATCTGTTCATTGAATATATC TTGGAC | GCTGCACGTGCTTTCTCTTCAAG CTC |
| 2. | ***β-actin*** | GCACCACACCTTCTACAATG | TGCTTGCTGATCCACATCTG |

**Table S5:** List of the primers used in BG4 ChIP:

| Sl. No. | Gene Name | Forward primer (5'- 3') | Reverse primer (5'- 3') |
|---|---|---|---|
| 1. | SMARCAL1 | AGGCGGACGAGACCAATGC | TCTTCCCCTCTAGGCAGGGAC |
| 2. | SMARCAL1 | CCCTGGGTGAGAAAGTGGTA | TCACCCAGCCTCAAGGTAAG |

| | | | |
|---|---|---|---|
| 3. | Non-G4 region | ATAGCTGCCCAGAGGCCTAA | CCGGAACATTTATTTTCCTATGAGC |

**Table S6:** LNA-FISH probe used in confocal microscopy:

| Sl. No. | Name | Sequence (5'-3') | Modification |
|---|---|---|---|
| 1. | Telomere antisense probe | CCCTAACCCTAACCCTAACCC | 5'-3'Tye 665 |

**Table S7:** Thermodynamic parameters of TMPyP4 binding to *SMARCAL1* P3 G-quadruplex as determined by ITC:

| System | Model | $K_d$ (M) | n | ΔH (kJ $mol^{-1}$) | ΔS (J $K^{-1}$ $mol^{-1}$) | ΔG (kJ $mol^{-1}$) |
|---|---|---|---|---|---|---|
| **P3** | Multiple sites | $K_d1=0.59 \times 10^{-6}$<br>$K_d2=0.81 \times 10^{-6}$ | n1=1.05<br>n2=1.94 | $\Delta H_1=60.02$<br>$\Delta H_2=-101.2$ | $\Delta S_1=320.5$<br>$\Delta S_2=-222.7$ | $\Delta G_1= -35.53$<br>$\Delta G_2= -34.81$ |

**Table S8:** Thermodynamic parameters of BRACO-19 binding to *SMARCAL1* P3 G-quadruplex as determined by ITC:

| System | Model | $K_d$ (M) | n | ΔH (kJ $mol^{-1}$) | ΔS (J $K^{-1}$ $mol^{-1}$) | ΔG (kJ $mol^{-1}$) |
|---|---|---|---|---|---|---|
| **P3** | Multiple sites | $K_d1=3.133 \times 10^{-8}$<br>$K_d2=1.000 \times 10^{-9}$ | n1=10.00<br>n2=4.830 | $\Delta H_1=-18.84$<br>$\Delta H_2=-21.21$ | $\Delta S_1=80.48$<br>$\Delta S_2=101.2$ | $\Delta G_1=-42.84$<br>$\Delta G_2= -51.38$ |

#### 3. Supporting discussions

**3.1. Thermodynamic profile of TMPyP4 and BRACO-19 with P3 G4:** To investigate the thermodynamics of the binding reactions, we performed ITC and evaluated the changes in enthalpy (ΔH), entropy (ΔS), Gibb's free energy (ΔG), and the equilibrium dissociation constant ($K_d$) (Table S7 and S8). The thermograms of P3 G4 with both TMPyP4 and BRACO-19 showed 'multiple-site' mode of binding. The negative value of Gibb's free energy (ΔG) suggested a forwarded, spontaneous formation of quadruplex-ligand complex (Figure S5A and Figure S5B). Consistent with the CD & ITC experiments, P3 G4 exhibited significant binding with TMPyP4 and BRACO-19. Stabilization of the G4 with TMPyP4 and BRACO-19 was also quite evident from the $K_d$ value and the rise in thermal stability of the P3 G4.

**3.2. *In cellulo* effect of TMPyP4 on the promoter G-quadruplex of *SMARCAL1*:** We also investigated the effect of TMPyP4 on *SMARCAL1* promoter activity by dual-luciferase assay with luciferase vector constructs pSMARCAL1-WT (with putative G-quadruplex motif) and pSMARCAL1-null (without G-quadruplex motif) (Figure S6A). We treated U2OS (Figure S6B) and Saos-2 (Figure S6C) cell lines with 10, 20 & 40 μM TMPyP4 for 24 h. The promoter activity of the wild type construct gradually decreased upon the increasing concentration of TMPyP4 in both the cell lines, compared to the control setup with no TMPyP4 treatment. But this reduction in the promoter activity was not observed in mutant construct without G-quadruplex forming sequence (Figure S6B & S6C). This dose-dependent mode of inhibition by TMPyP4 can surely suggest that TMPyP4 interacted with the P3 G4 in the promoter region, thus stabilizing it and reducing the transcription rate of *SMARCAL1*.

**3.3. Downregulation of SMARCAL1 induces DNA damage throughout the genome:** To evaluate if the SMARCAL1 downregulation was coupled with the increase in DNA damage at genomic level, the presence of p-γH2AX, a novel, potent biomarker for DNA double stranded breaks[2], was also evaluated. Both U2OS and Saos-2 displayed a significant rise in the p-γH2AX immunofluorescence thus DNA damage at whole genomic level (Figure S11A, S11B, S11C and

S11D). U2OS accumulated much higher DNA damage due to increased TMPyP4. Even BRACO-19 treatment also induced significant DNA damage in U2OS cells, which can be clearly observed by the rise of p-γH2AX foci in those cells. No significant DNA damage was observed in the untreated ALT cells. By silencing *SMARCAL1* gene by siSMARCAL1, we also observed a similar rise in DNA damage. Thus, it can be stated that TMPyP4 and BRACO-19 mediated stabilization of *SMARCAL1* promoter G4 surely have increased the global DNA damage and replication stress significantly at telomeres of ALT positive cells, which clearly supports the stress regulatory role of SMARCAL1.

#### 4. Supplementary Figures and Figure legends

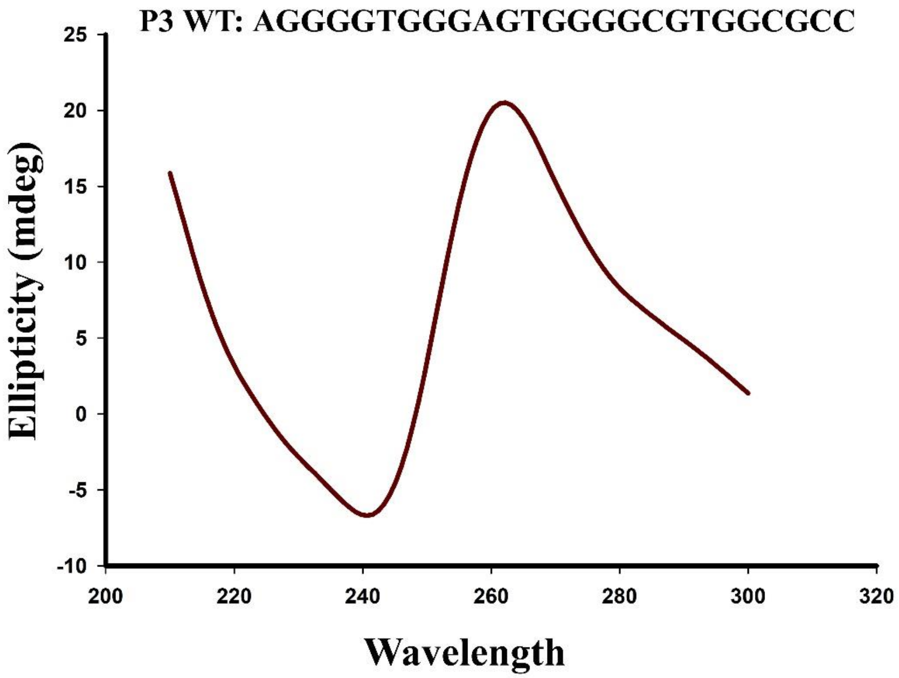


**Figure S1.** CD Spectra of the wild type oligonucleotide sequence (P3 wt), annealed in 10 mM potassium phosphate buffer with 100 mM potassium chloride.

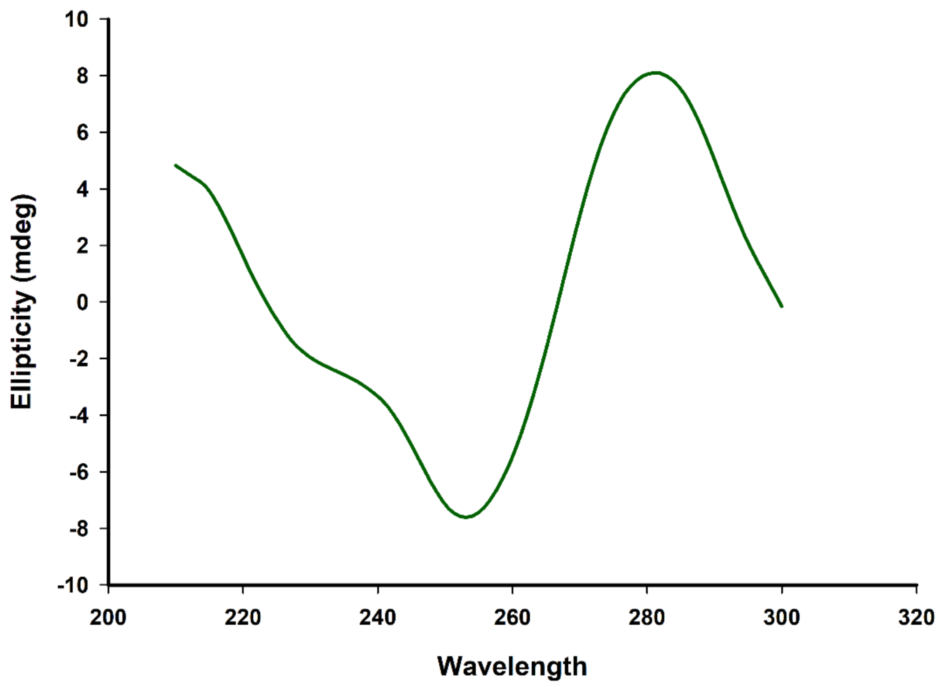


**Figure S2.** CD Spectra of the mutated oligonucleotide sequence (P3 mut), annealed in 10 mM potassium phosphate buffer with 100 mM potassium chloride.

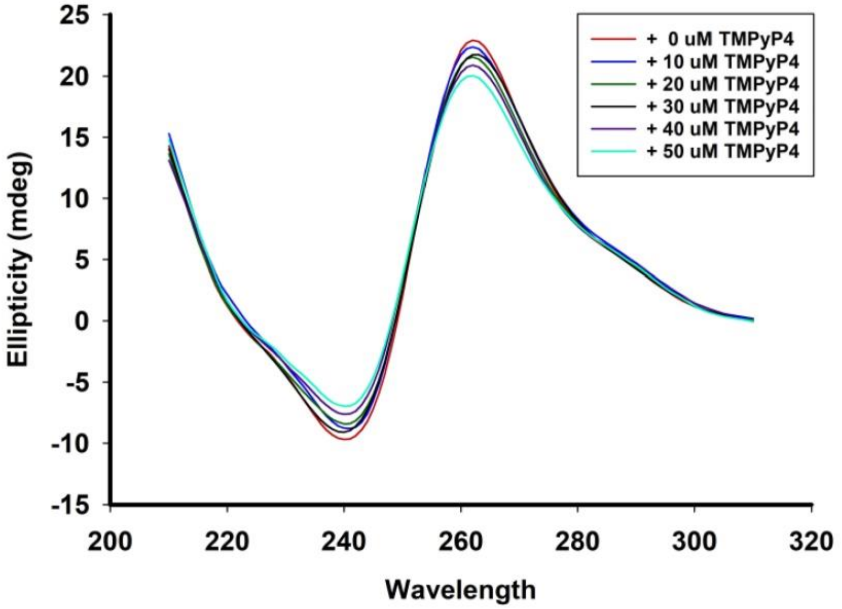


**Figure S3. CD spectra of P3 wt due to TMPyP4 interaction.** Study of the interaction of the P3 wt sequence titrating with TMPyP4 using CD spectroscopy. CD spectra of the P3 wt sequence titrated with increasing concentration of TMPyP4, suggesting no change in the nature of the CD spectra of the G4 forming oligonucleotides upon titration.

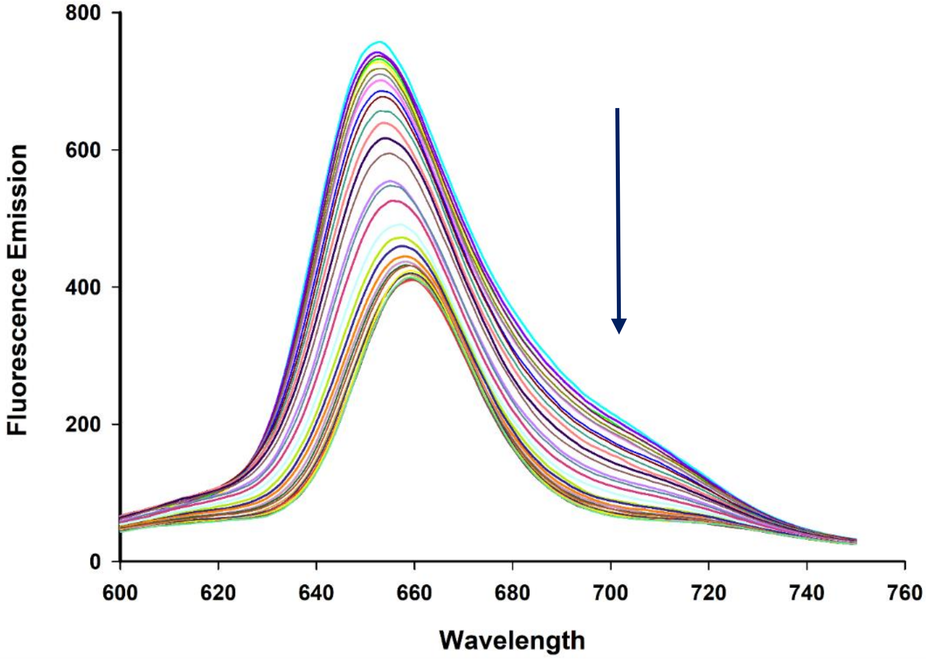


**Figure S4.** Fluorescence emission spectra of TMPyP4 titrated with increasing concentration of G4 forming sequence (P3 wt) till saturation.

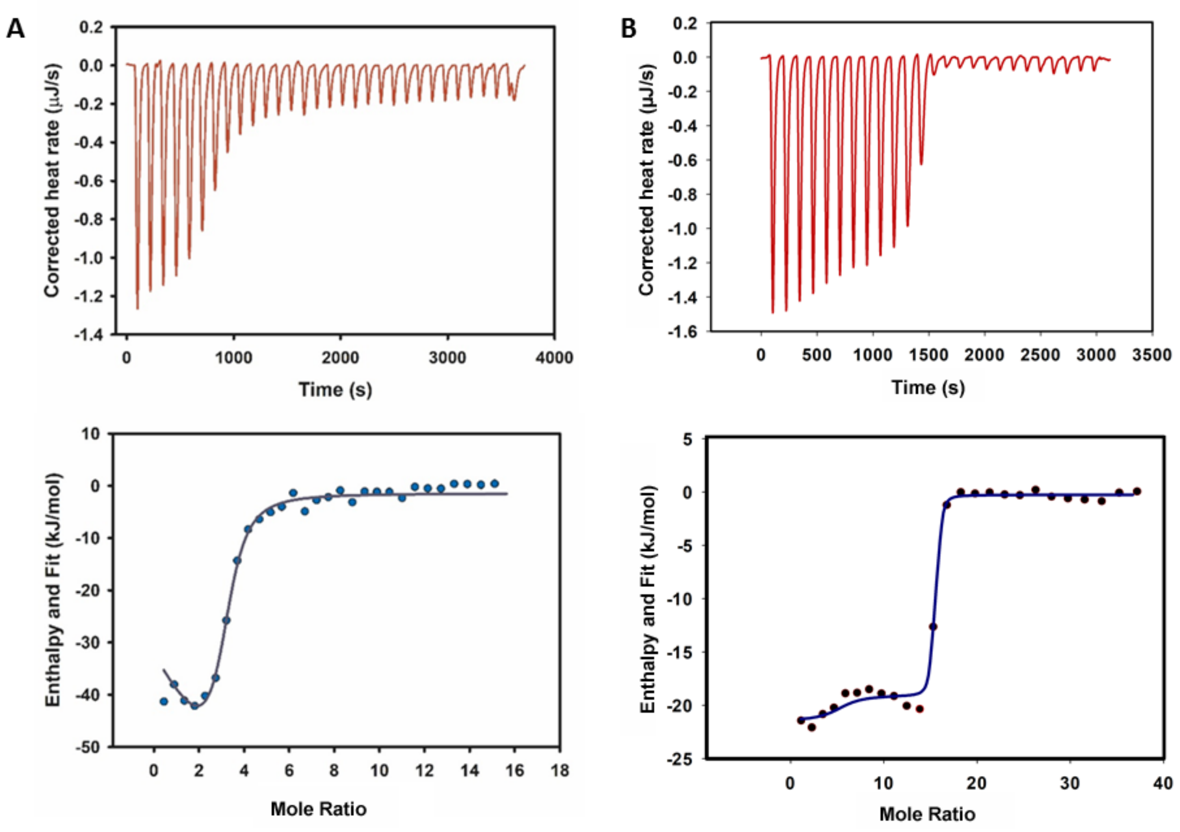


**Figure S5. ITC profiles of P3 wt with TMPyP4 and BRACO-19.** ITC experiments were performed with the P3 wt oligonucleotides (cell) titrated with increasing concentration of (A) TMPyP4 (syringe) and (B) BRACO-19 (syringe) in different experiments. Then, the profile of each complex was shown. The upper panel represented the corrected heat rate with every injection leading towards saturation. The lower panel dictated the curve fit of the corrected heat rate profiles to elucidate the mode of binding for quantification of the binding profiles.

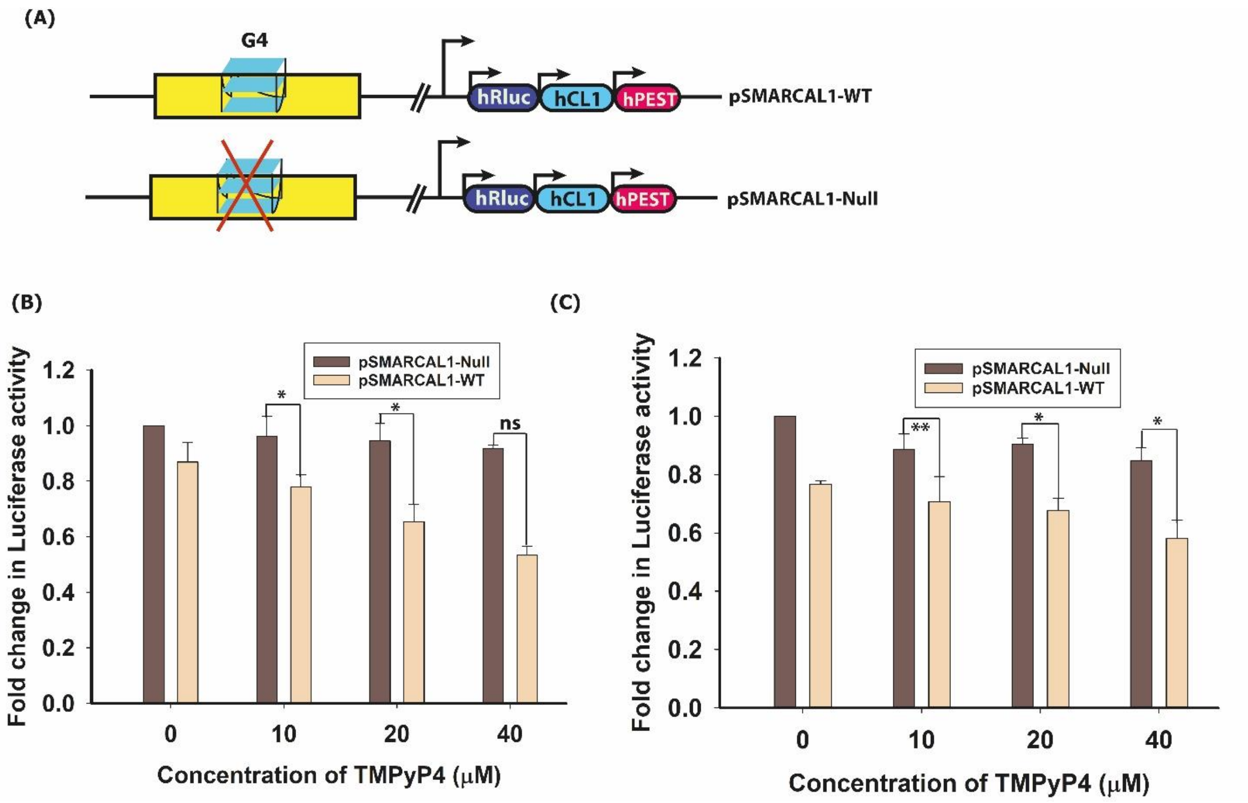


**Figure S6. Dual-luciferase assay** (A) The schematic representation of the luciferase vector and the constructs containing human *SMARCAL1* genomic region containing predicted G4 motif in wild type and mutated form. The dual luciferase assay was performed with pSMARCAL1-WT and pSMARCAL1-Null constructs in U2OS (B) and Saos2 (C) cells in presence (10, 20 and 40 μM) and absence of TMPyP4. The relative luciferase activity of each plasmid was calculated after normalizing *Renilla* luciferase with firefly luciferase activity and represented as fold change.

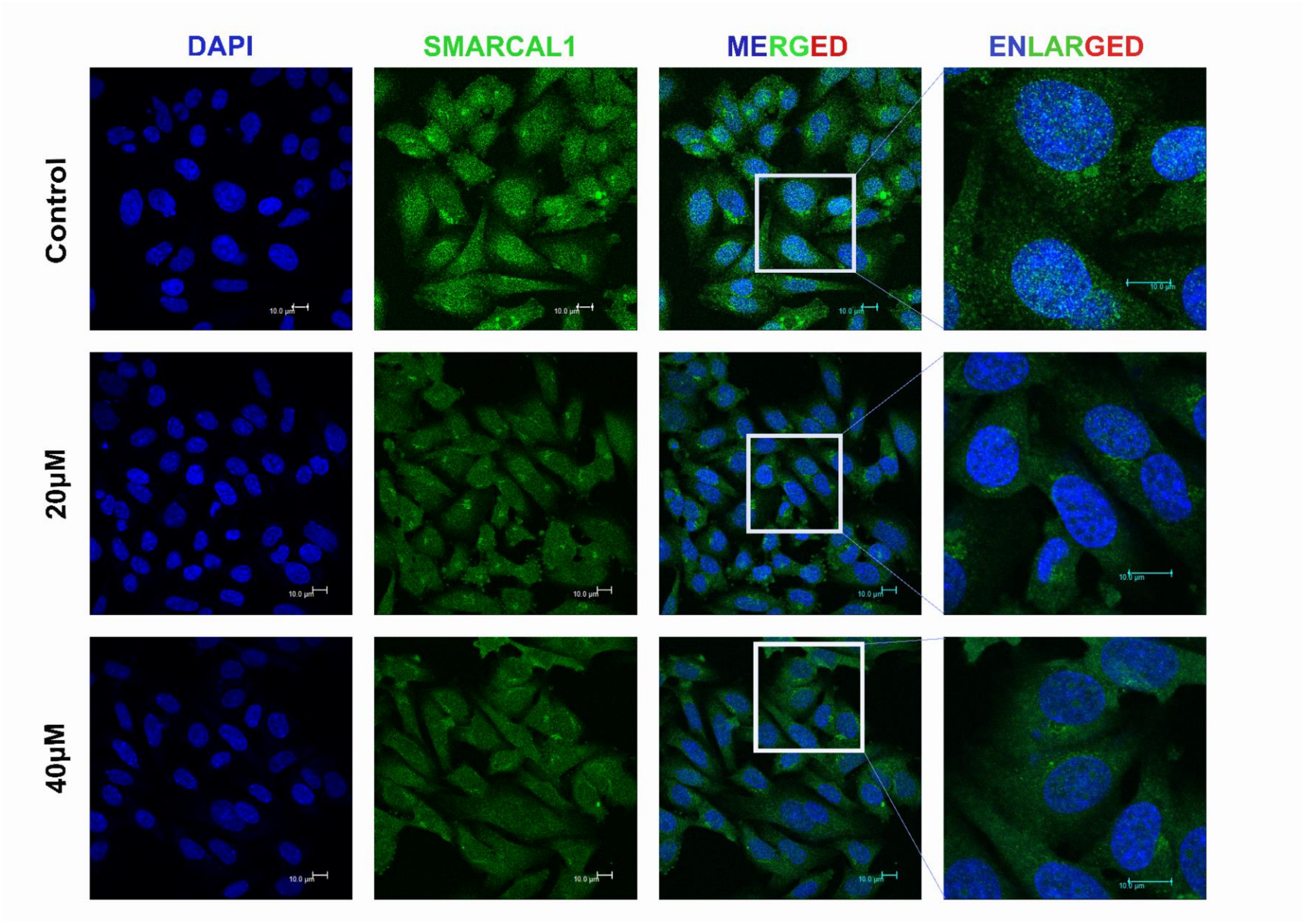


**Figure S7. Immunocytochemistry of SMARCAL1**. Immunostaining with antibody against SMARCAL1 (green) was done in Saos2 cell line, treated with 20 and 40 µM TMPyP4 for 24 h and compared with that of control. The nuclei stained with DAPI (blue) and merged images of SMARCAL1 and DAPI were also shown. Scale bar: 10 µm.

A

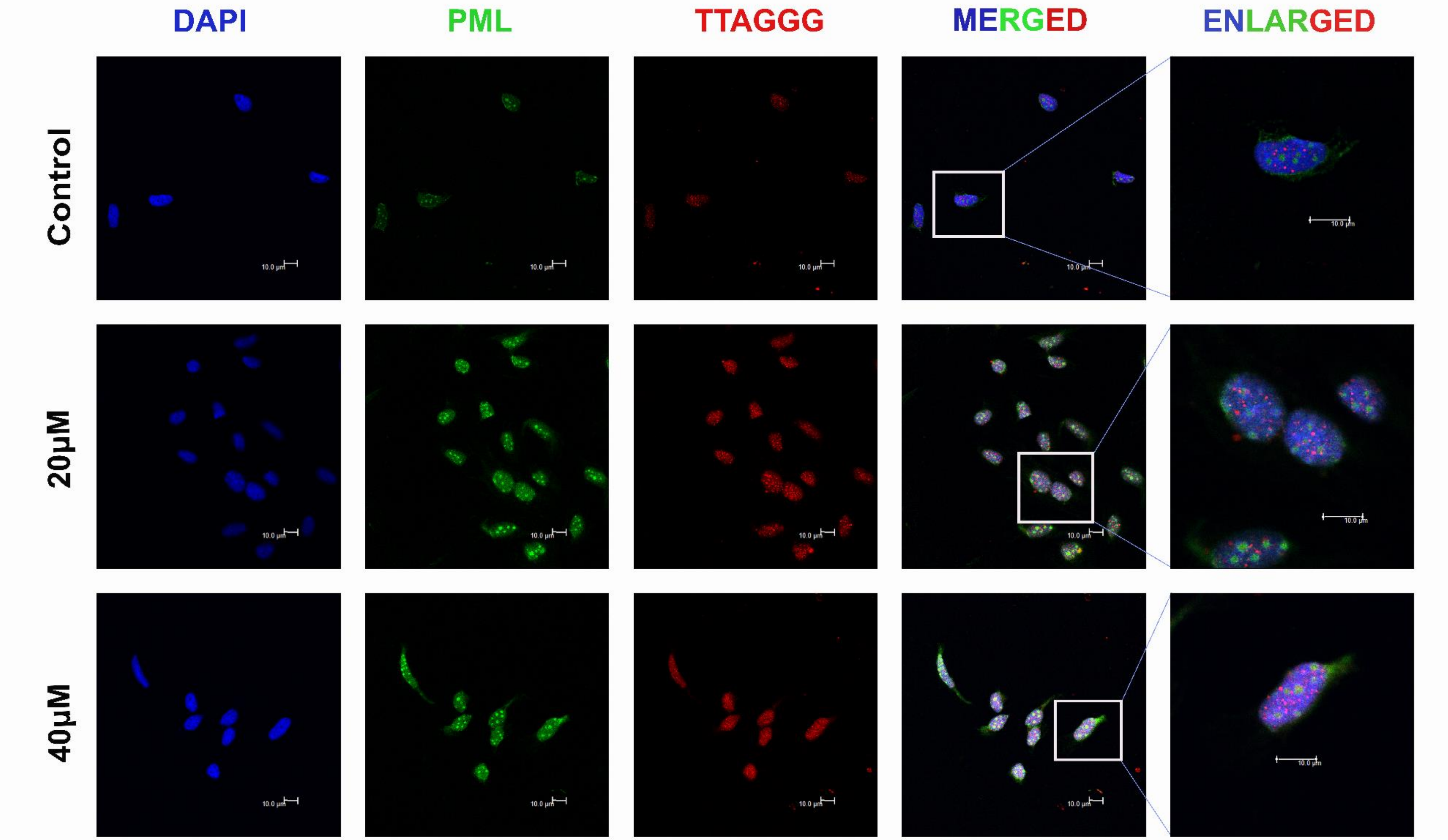

**B**

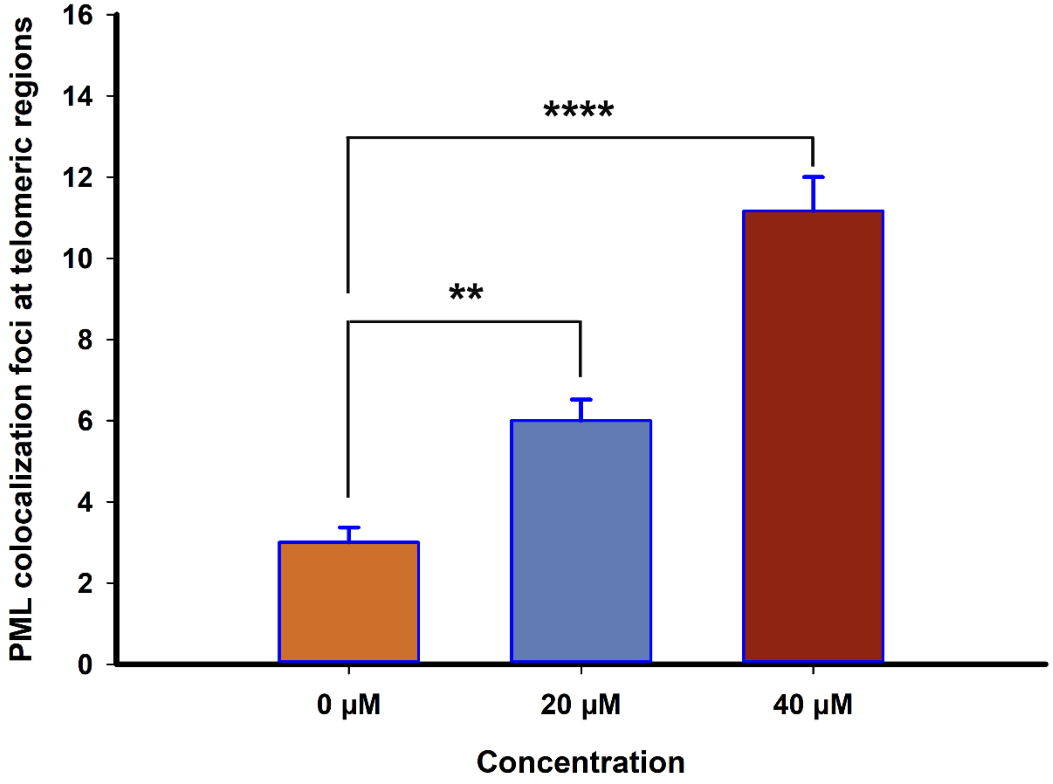


**Figure S8. Recruitment of PML bodies (APB foci) at ALT telomeres due to SMARCAL1 downregulation**. (A) Cells were treated with 20 and 40 µM TMPyP4 for 24 h and compared with that of control. Combined immunofluorescence and fluorescence in situ hybridization (FISH) analyses the colocalization of PML and telomeres (TTAGGG) in Saos2 cells. The nuclei stained with DAPI (blue) and merged images of PML (green) and telomere (red) were shown. Representative images were shown. Scale bar: 10 µm. (B) Graphical representation showing the numbers of PML colocalization foci at telomeric regions. p-values are calculated in comparison with control (0 µM) dataset.

A

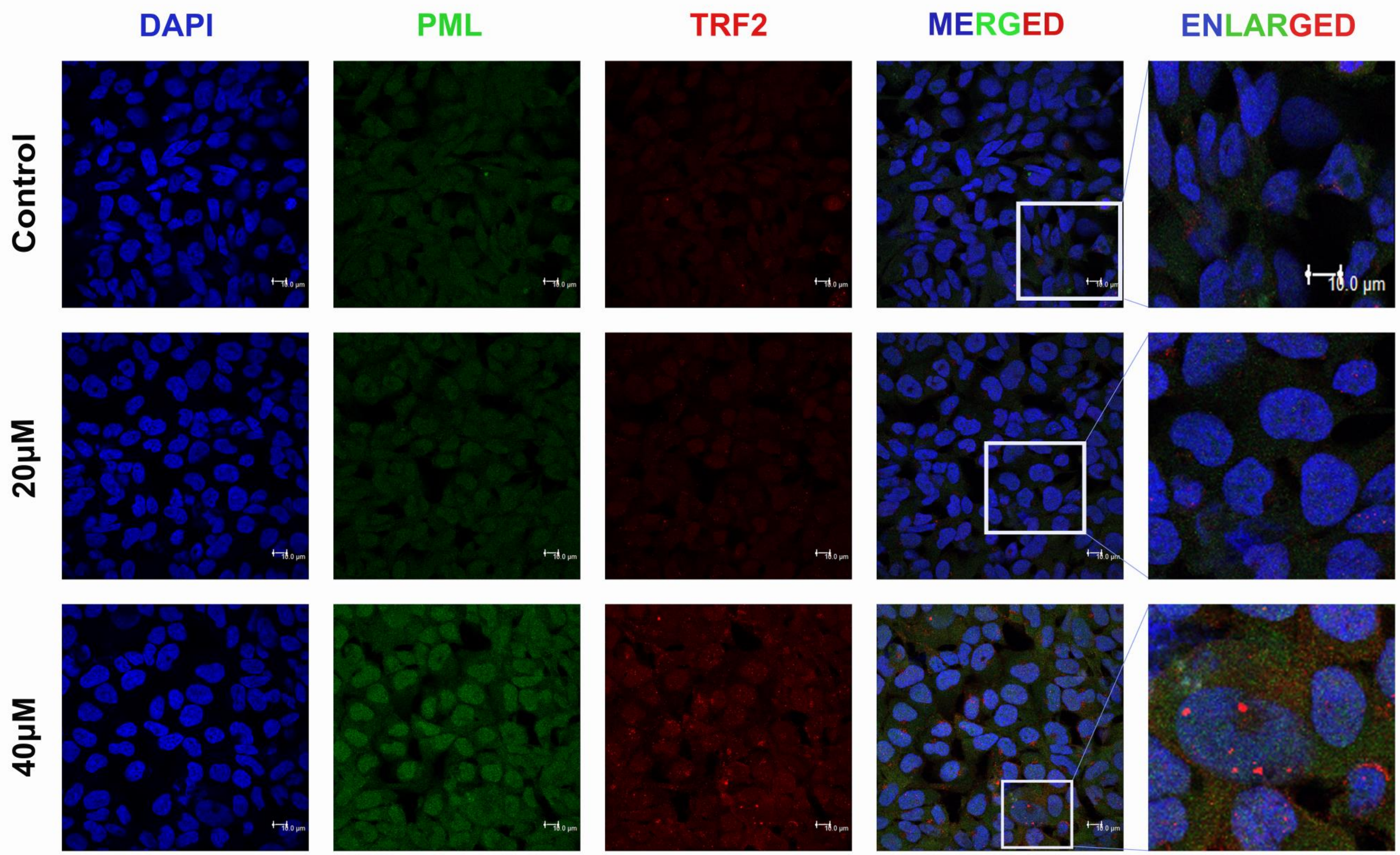

**B**

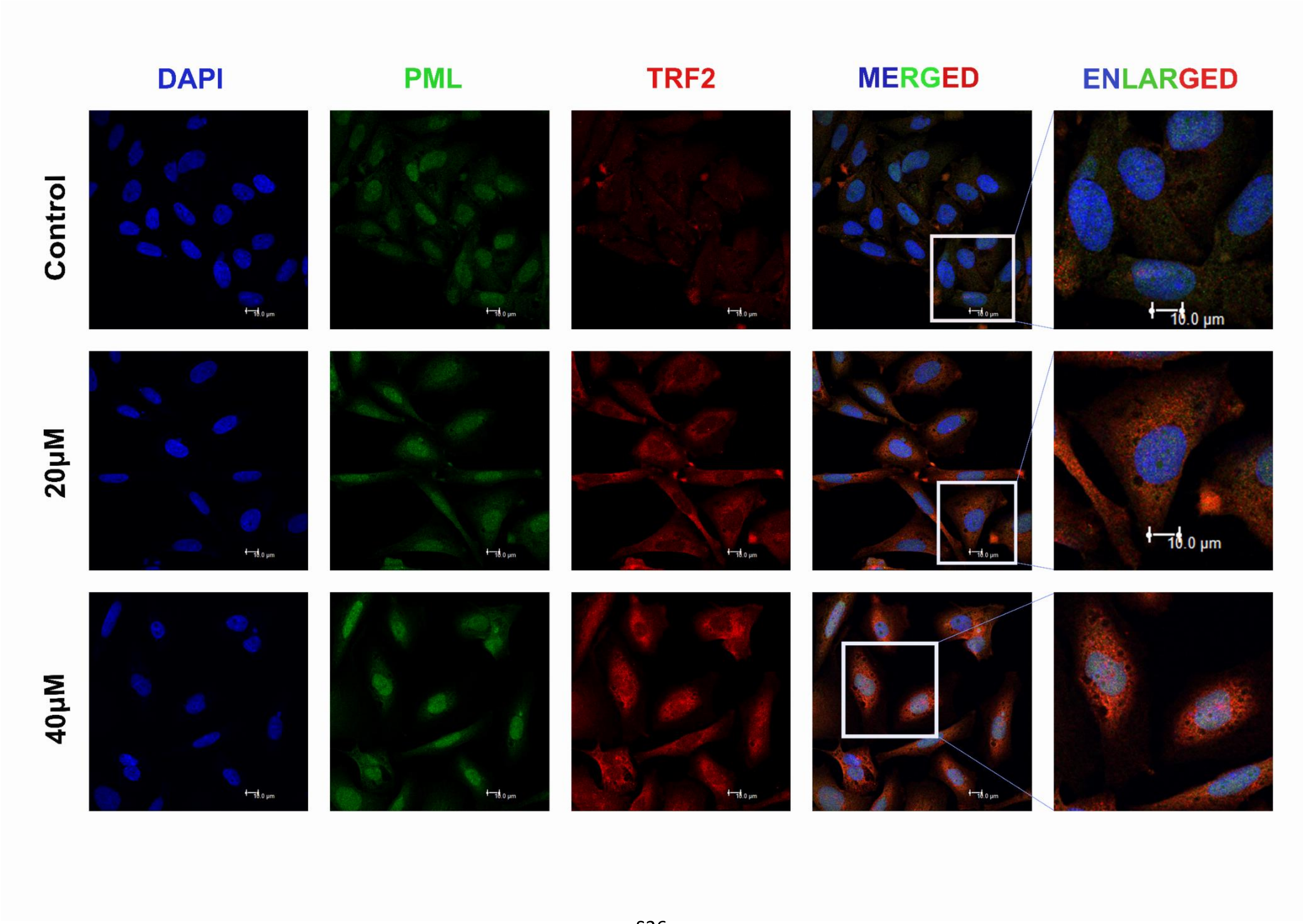
DAPI
PML
TRF2
MERGED
ENLARGED
Control
20μM
40μM
10.0 μm
10.0 μm

**Figure S9. Interaction profile and co-localization of PML and TRF2 protein.** (A) U2OS and (B) Saos-2 Cells were treated with 20 and 40 μM TMPyP4 for 24 h and compared with that of control. Immunofluorescence of PML (green), TRF2 (red) and their colocalization were analyzed in Saos2 cells. The nuclei stained with DAPI (blue) and merged images of PML (green) and TRF2 (red) were shown. Representative images were shown. Scale bar: 10 μm.

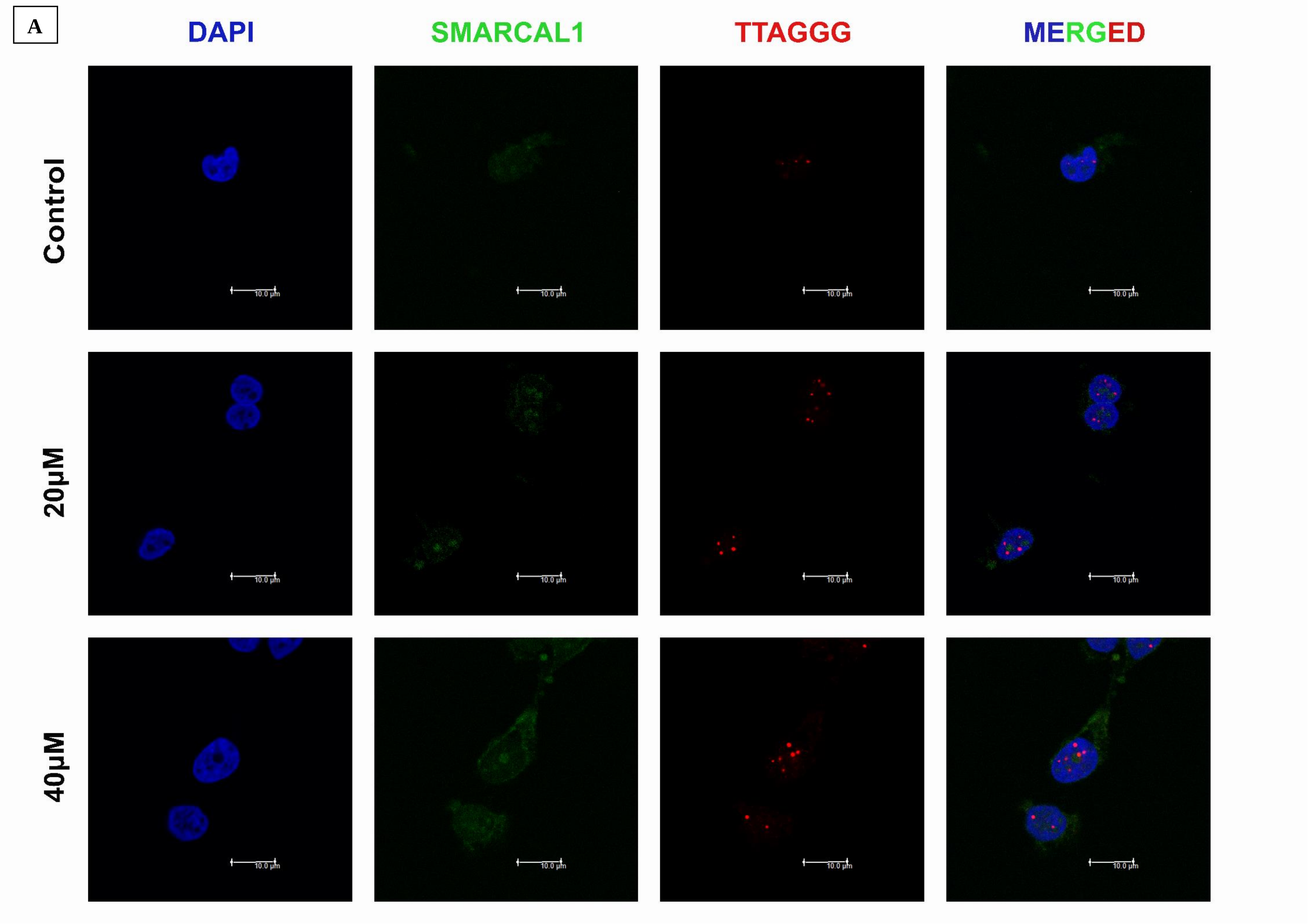
A
DAPI
SMARCAL1
TTAGGG
MERGED
Control
20μM
40μM
10.0 μm

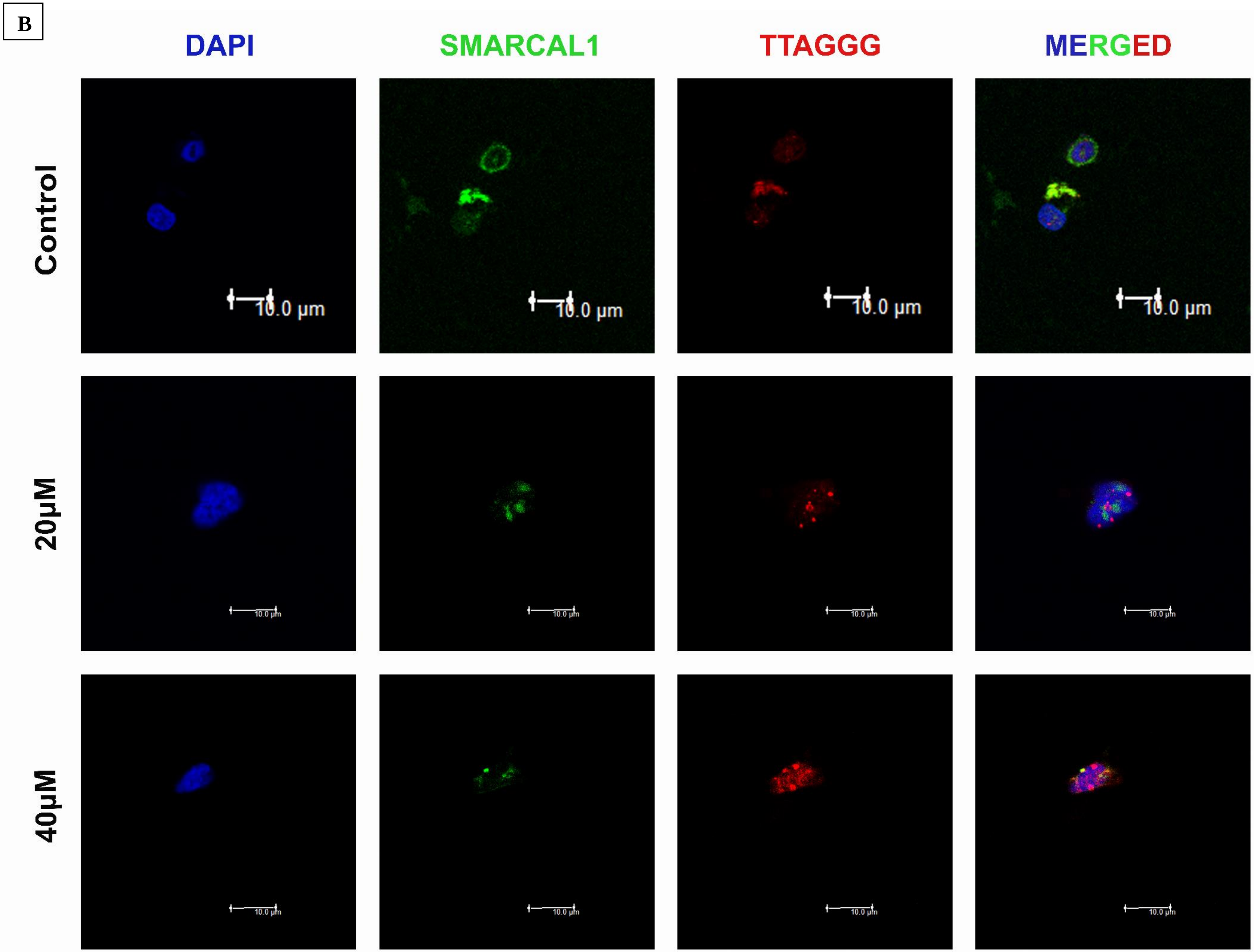
B
DAPI
SMARCAL1
TTAGGG
MERGED
Control
20μM
40μM
10.0 μm

**Figure S10. Loss of SMARCAL1 accumulation in ALT telomeres due to TMPyP4 mediated stabilization of the *SMARCAL1* G-quadruplex.** Cells were treated with 20 and 40 μM TMPyP4 for 24 h and compared with that of control. Combined immunofluorescence and fluorescence in situ hybridization (FISH) analyses the removal of SMARCAL1 from telomeres (TTAGGG) in U2OS (A) and Saos2 (B). The nuclei stained with DAPI (blue) and merged images of SMARCAL1 (green) and telomere (red) were shown. Representative images were shown. Scale bar: 10 μm.

**A**

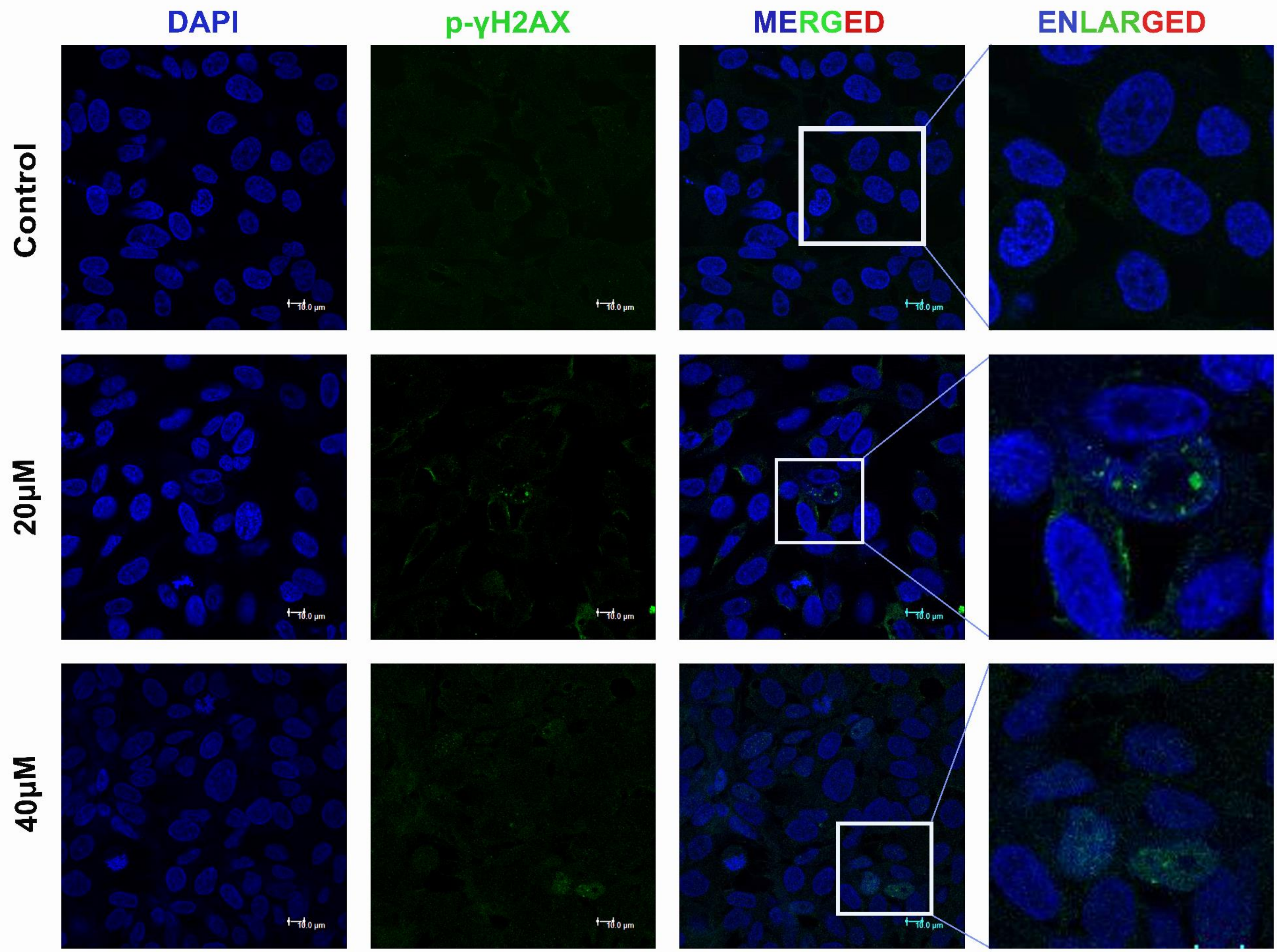

B

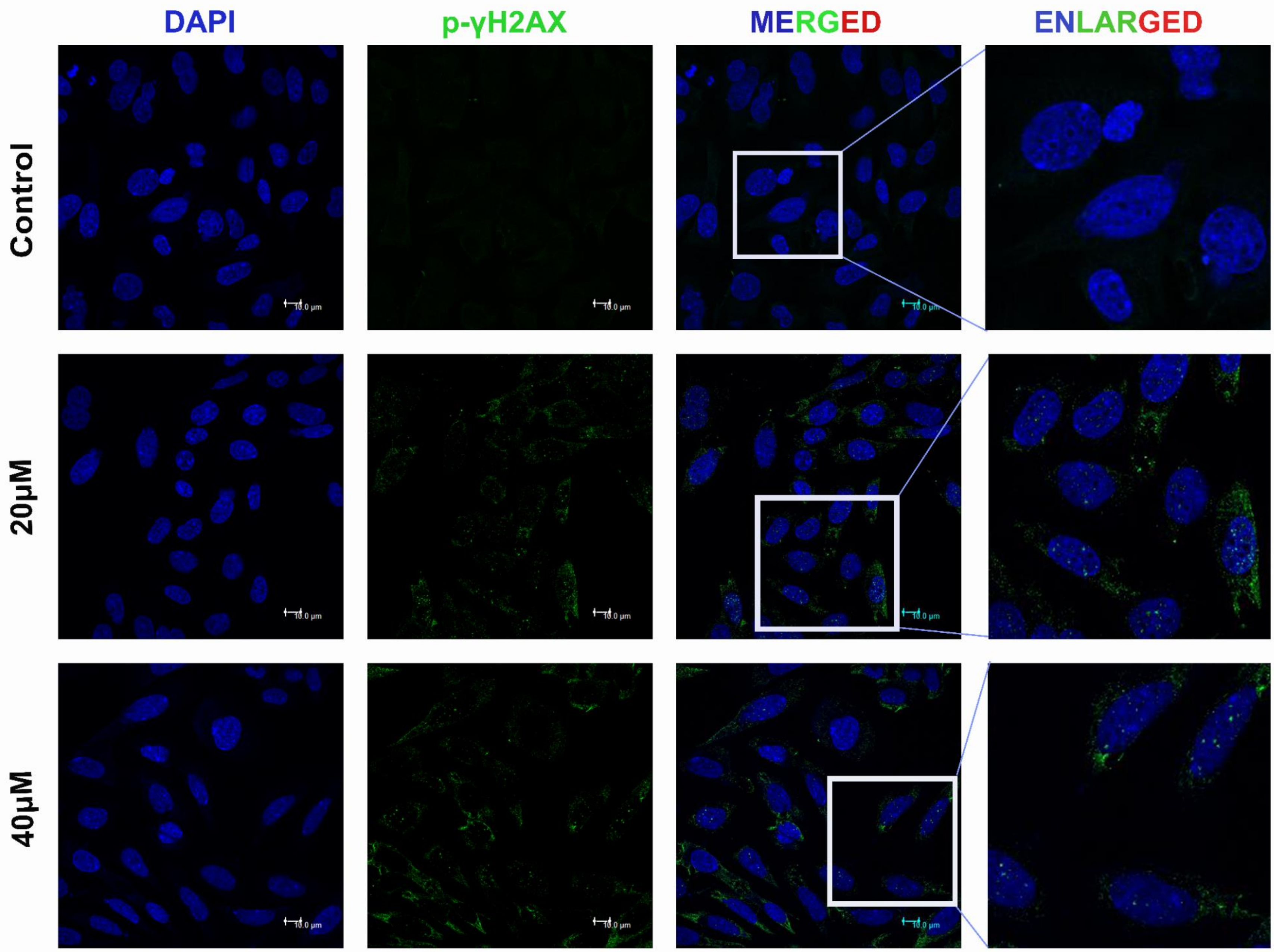

C

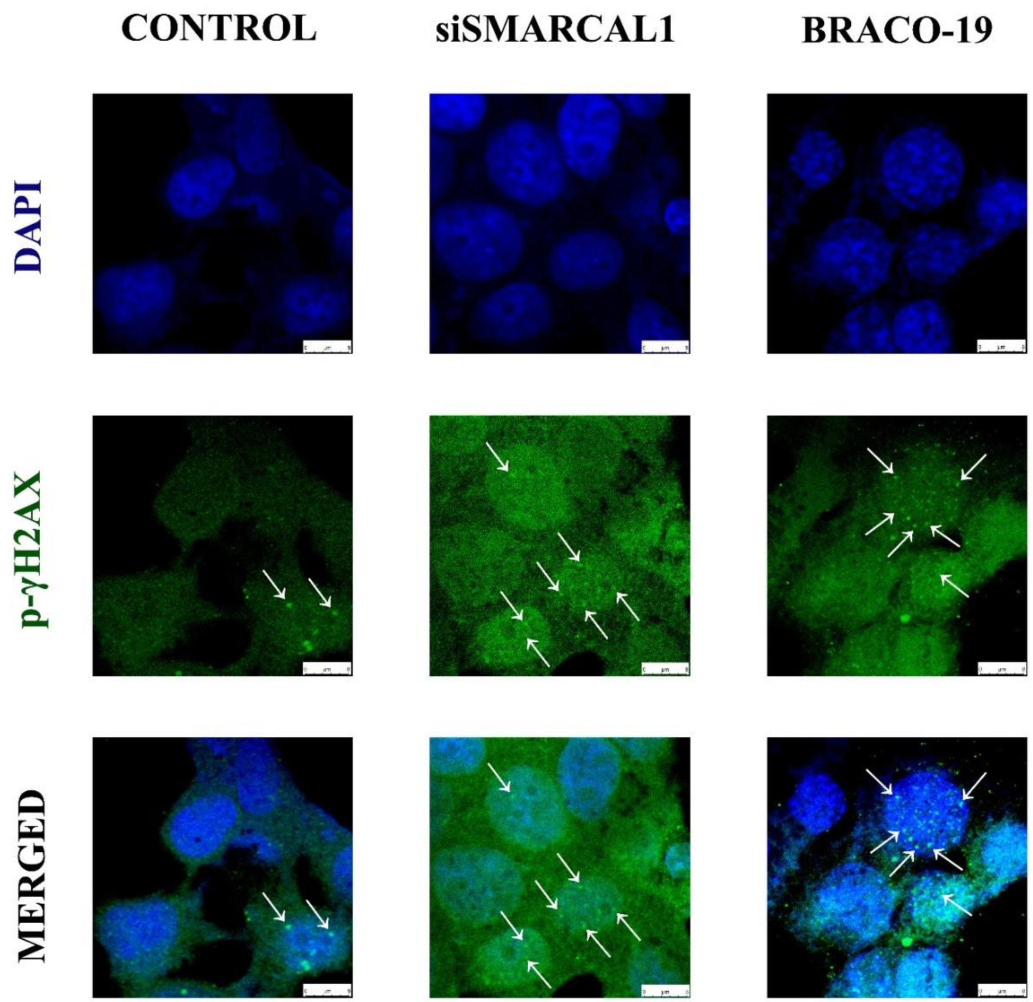


D

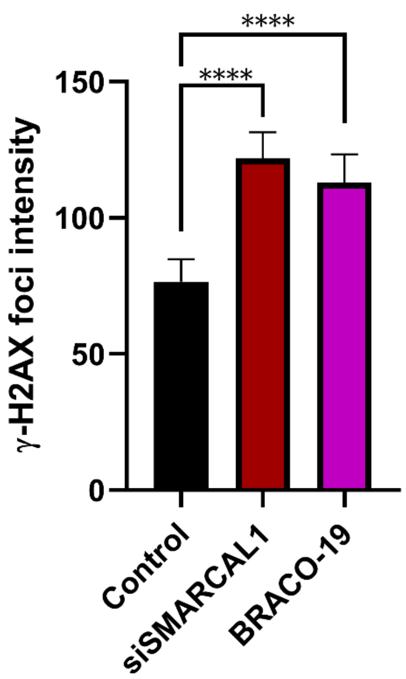


**Figure S11. Loss of SMARCAL1 induces DNA damage at genomic level.** Cells were treated with 20 and 40 μM TMPyP4 for 24 h and compared with that of control. Immunofluorescence of p-γ H2AX (green) was analyzed. The nuclei stained with DAPI (blue) and merged images of p-γ H2AX (green) were shown in U2OS (A) and Saos2 (B). U2OS cells were also treated with 15 μM BRACO-19 and compared with siSMARCAL1 treatment (C). In both the cases, number of p-γ H2AX foci increases, reflecting the global DNA damage due to the downregulation of SMARCAL1 through stabilizing the SMARCAL1 G4 by TMPyP4 and BRACO-19. (D) Graphical representation of γ-H2AX fluorescence intensity in Control, siSMARCAL1 and BRACO-19 treated U2OS cells.

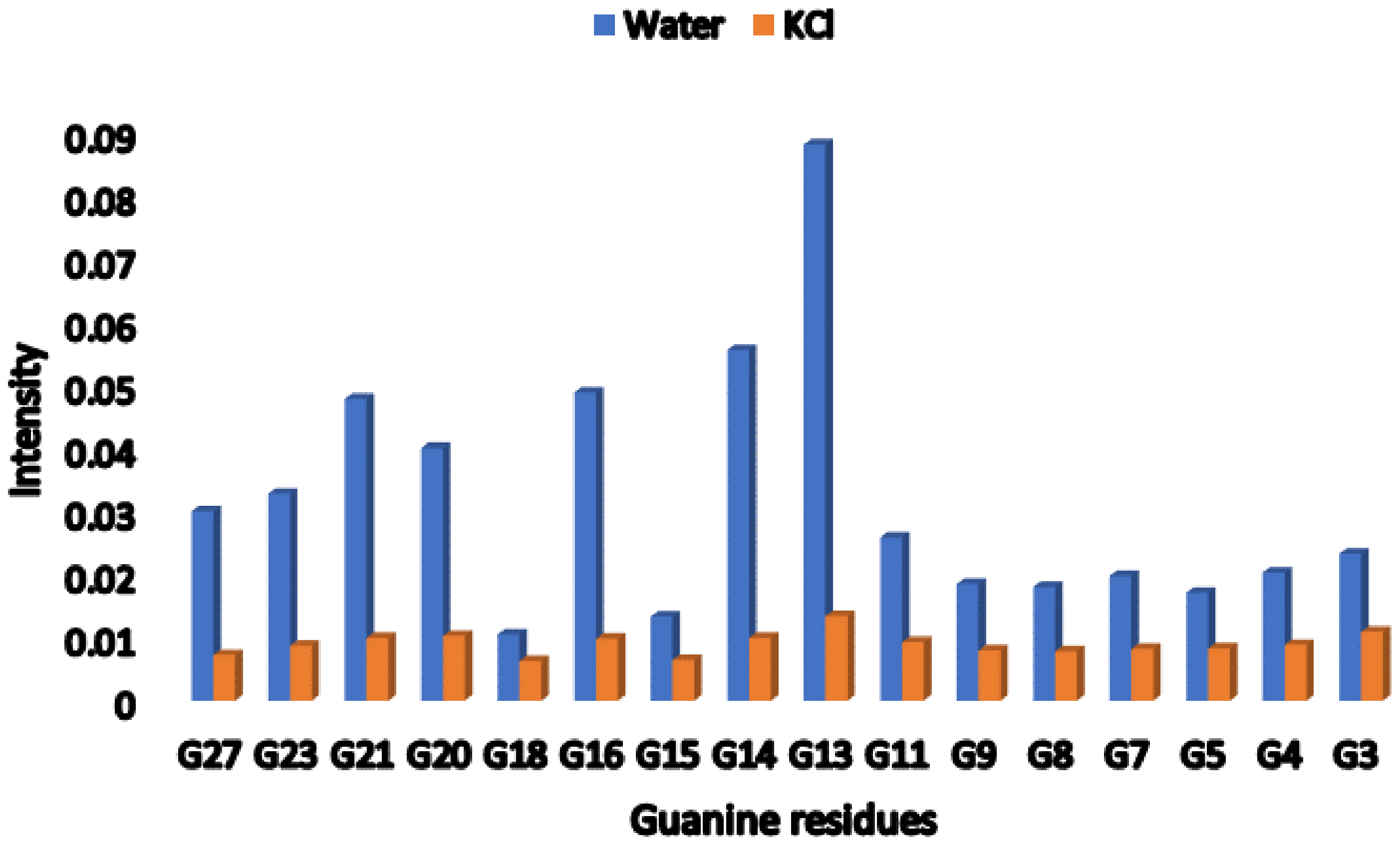


**Figure S12. DMS footprinting densitometric plot.** Intesity of all the bands (Guanine residues) were analysed and represented graphically.

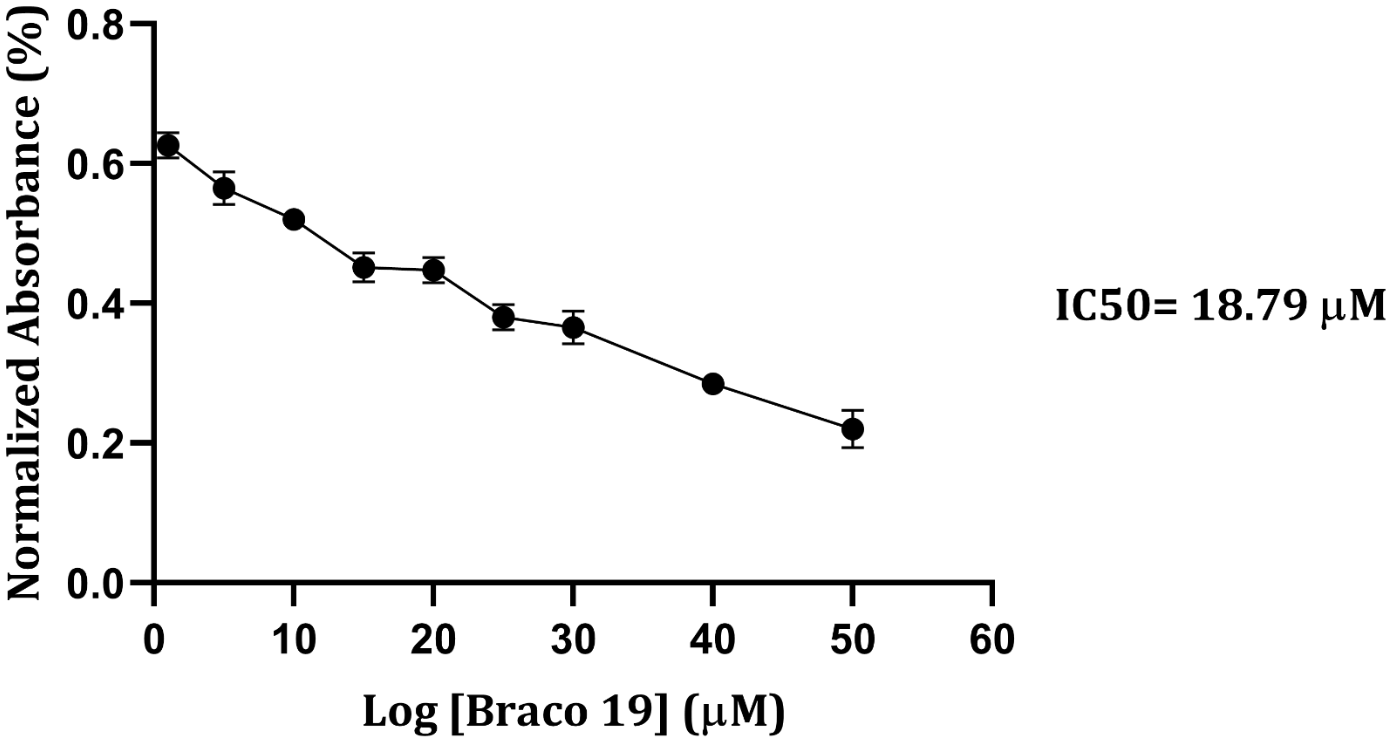


**Figure S13. MTT assay for BRACO-19 in U2OS cells.**

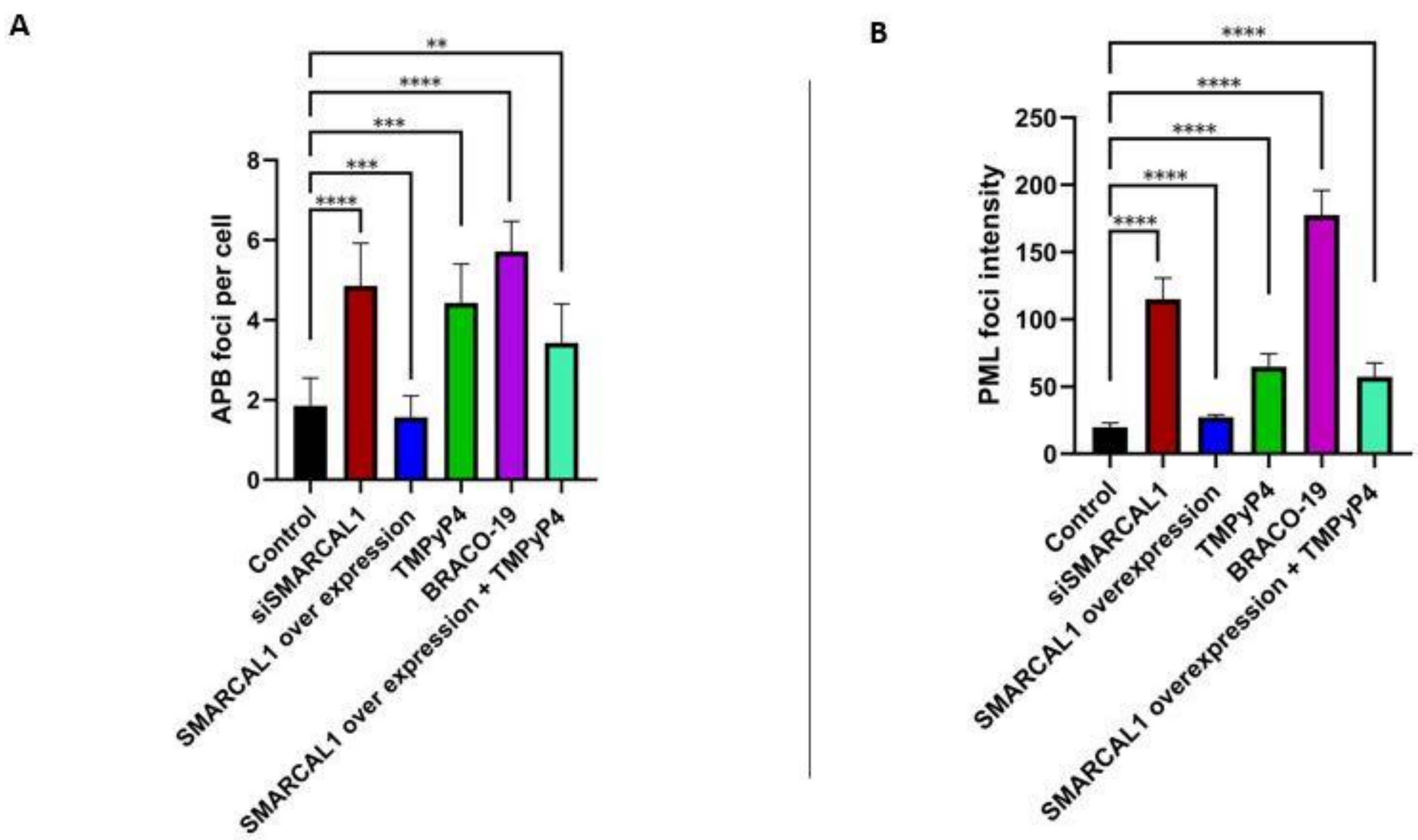


**Figure S14. Graphical representation of the APBs formation in different experimental sets.** (A) APB foci per cell and (B) PML foci fluorescence intensity has been quantified in U2OS cells (Figure 3) and represented graphically. p-values are calculated in comparison with control dataset.

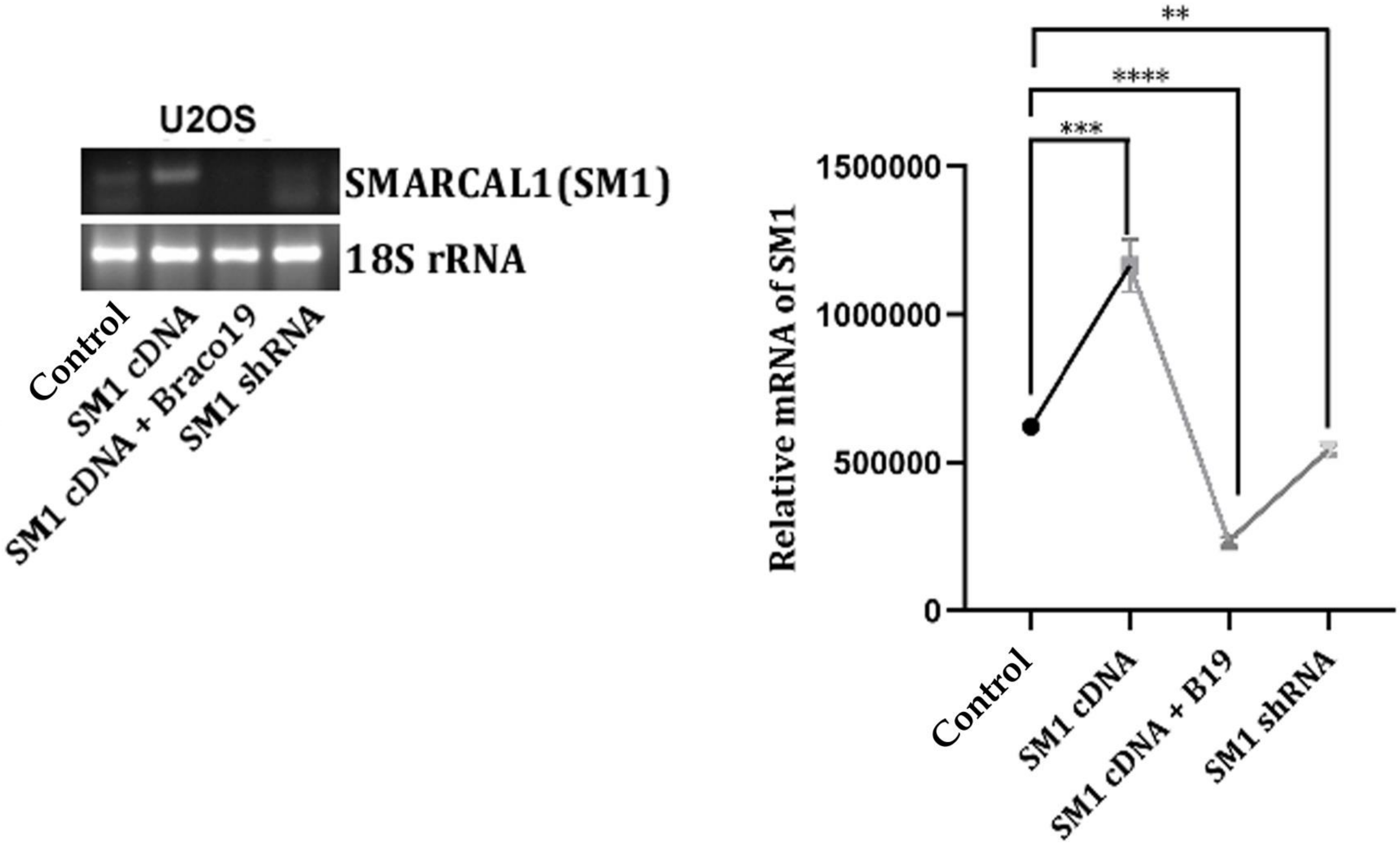


**Figure S15. Expression profile of SMARCAL1 mRNA in U2OS cells.** Analysis of SMARCAL1 expression profile in control, SMARCAL1 silencing, SMARCAL1 overexpression and BRACO-19 (B19) treated conditions. Relative mRNA expression was also represented graphically in comparison with 18S rRNA.

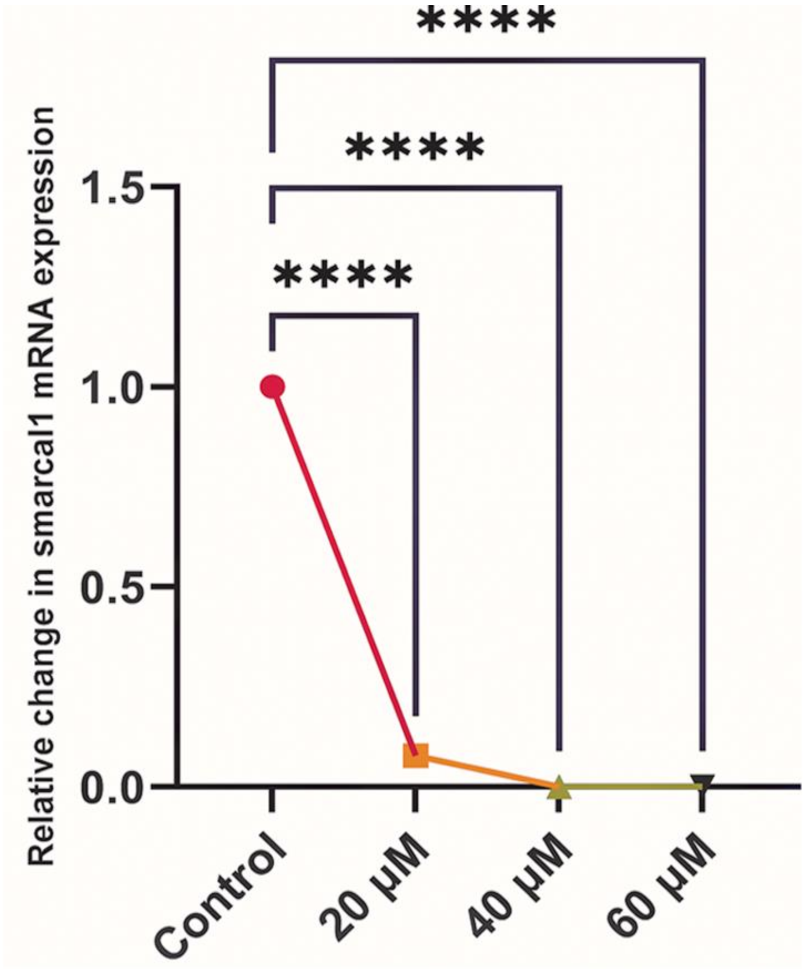


**Figure S16. Graphical analysis of SMARCAL1 mRNA expression profile in Saos-2 cells.** SMARCAL1 mRNA expression was checked by PCR in Saos-2 cells treated with TMPyP4 for 24 hrs.

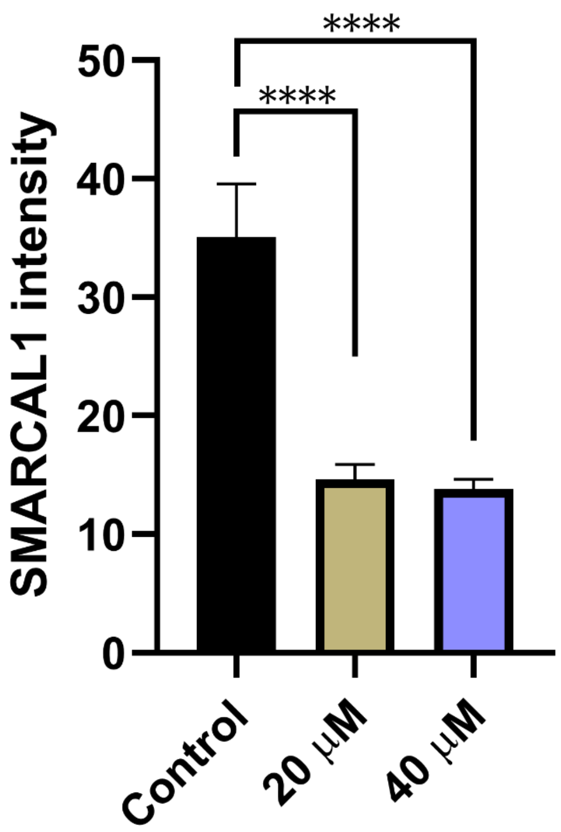


**Figure S17. Graphical representation of the fluorescence intensity of SMARCAL1 protein.** SMARCAL1 intensity is calculated from the nuclear region of U2OS cells in Control, 20μM and 40 μM TMPyP4 treatment and represented graphically. p-values are calculated in comparison with control dataset.

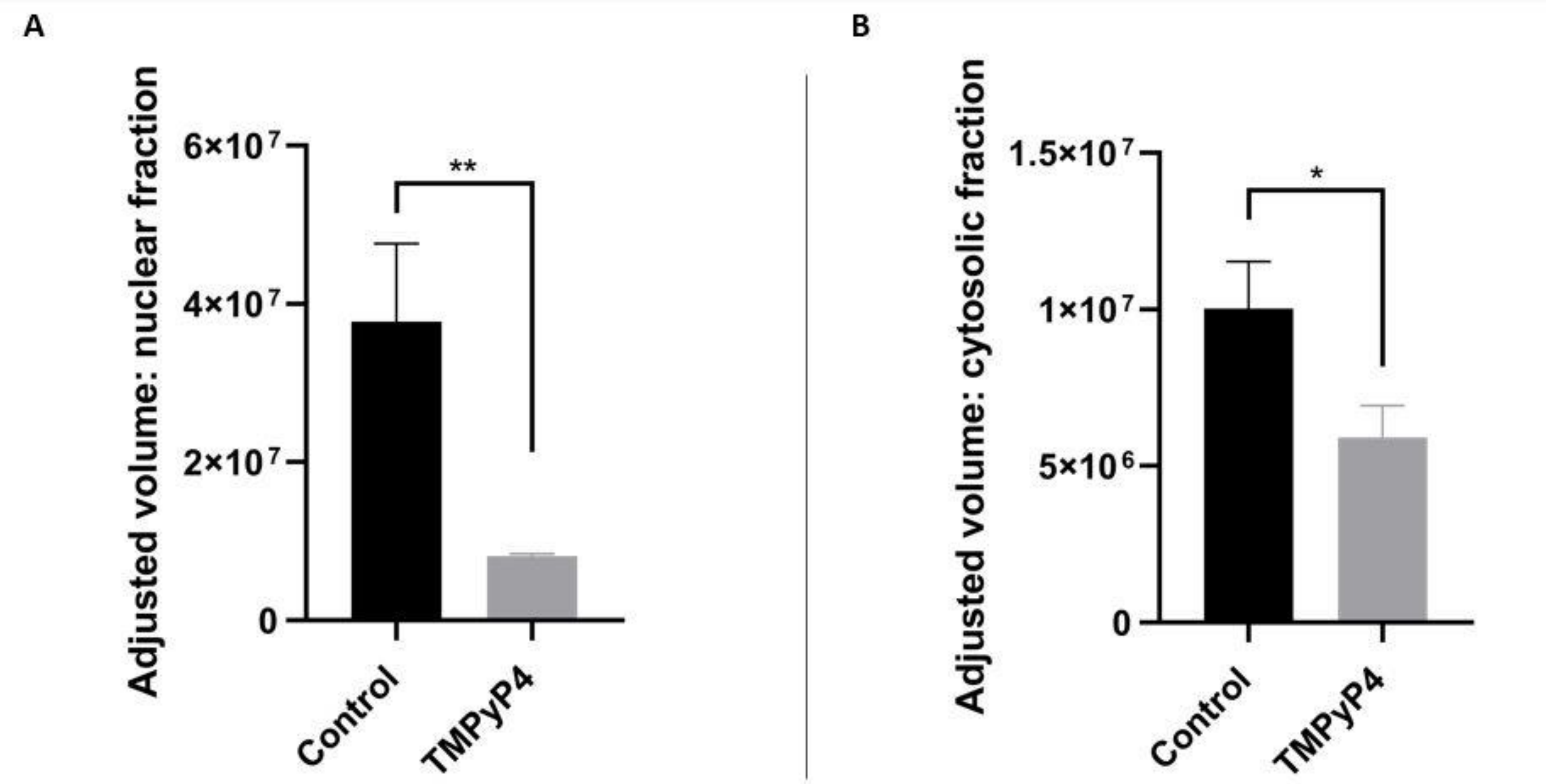


**Figure S18. Densitometric analysis of the Western Blot.** Densitometric values were plotted for both the nuclear (A) and cytosolic (B) fraction of SMARCAL1 protein in the absence and presence of TMPyP4 (40μM) in U2OS cells.
